# Watching Death in the Gerbil Cochlea Using Optical Coherence Tomography

**DOI:** 10.1101/2021.04.29.442005

**Authors:** Nam Hyun Cho, Haobing Wang, Sunil Puria

## Abstract

Because it is difficult to directly observe the morphology of the living cochlea, our ability to infer the mechanical functioning of the living ear has been limited. Nearly all of our knowledge about cochlear morphology comes from postmortem tissue that was fixed and processed using procedures that possibly distort the structures and fluid spaces of the organ of Corti. In this study, optical coherence tomography was employed to obtain in vivo and postmortem micron-scale volumetric images of the high-frequency hook region of the gerbil cochlea through the round-window membrane. The anatomical structures and fluid spaces of the organ of Corti were segmented and quantified in vivo and over a 90-minute postmortem period. The results show that some aspects of the organ of Corti are significantly altered over the course of death, such as the volumes of the fluid spaces, whereas the dimensions of other features change very little. We postulate that the fluid space of the outer tunnel and its surrounding tectal cells form a resonant structure that can affect the motion of the reticular lamina and thereby have a profound effect on outer-hair-cell transduction and thus cochlear amplification. In addition, the in vivo fluid pressure of the inner spiral sulcus is postulated to effectively inflate the connected sub-tectorial gap between the tectorial membrane and the reticular lamina. This gap height decreases after death, which is hypothesized to reduce and disrupt hair-cell transduction

## I. Introduction

A touchstone of biological research is the notion that structure determines function. In the auditory system, hearing function is closely associated with the intricate structure of the organ of Corti (OoC), the sensory epithelium within the spiral-shaped, fluid-filled cochlea of the inner ear. The OoC itself is attached to the basilar membrane (BM), which partitions the cochlear fluid space as it runs along the length of the spiral from the high-frequency basal hook region (Fig. 1a, b) to the low-frequency apical end of the cochlea.

**Fig. 1.**
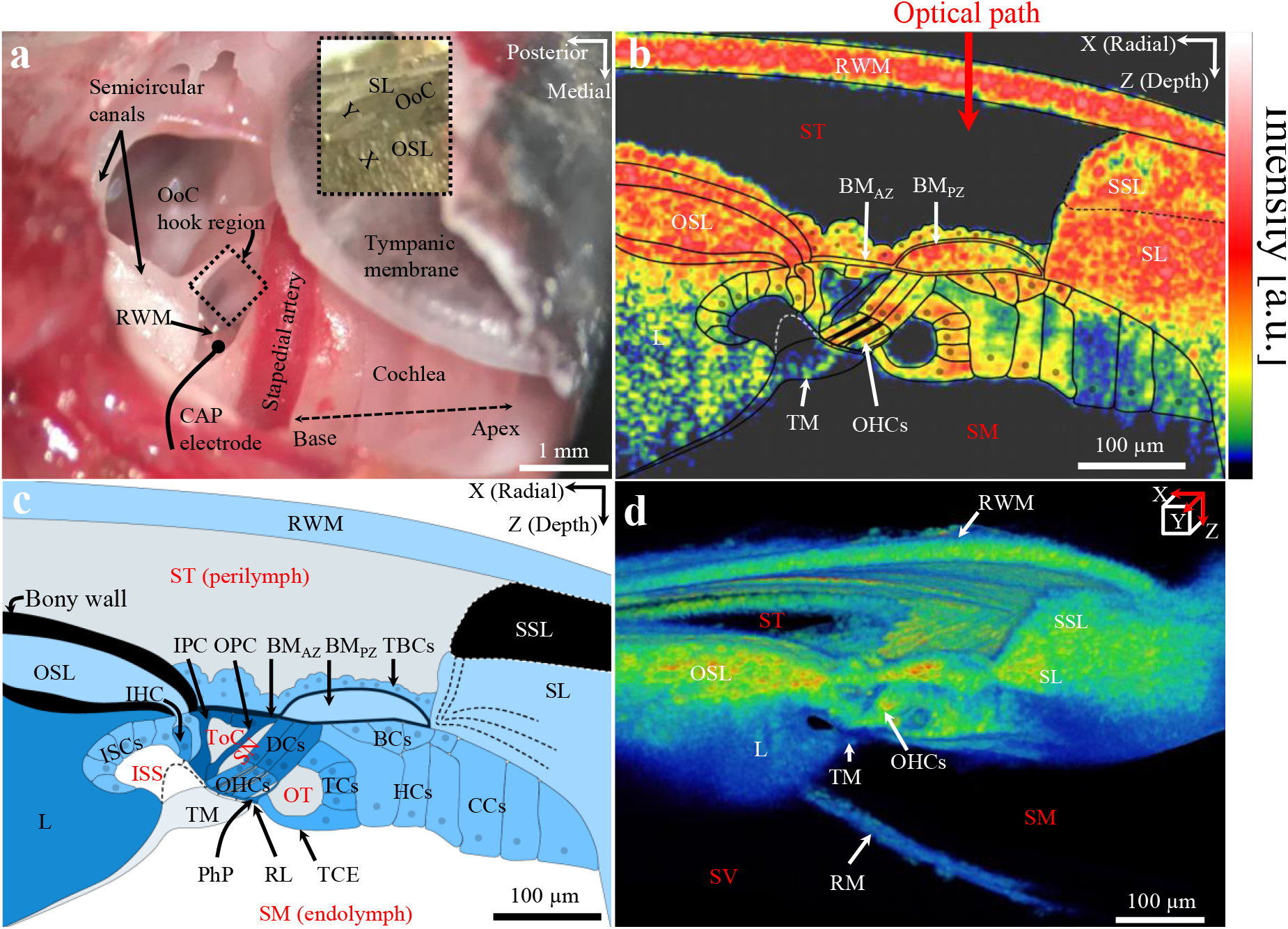
The anatomy of the gerbil ear, as depicted by multiple modalities. **a:** A photograph of the surgically opened right middle ear shows the region of the round-window membrane (RWM; black dotted square), through which the organ of Corti (OoC) in the “hook region” of the cochlea (inset image) can be imaged using optical coherence tomography (OCT). The inset image, the X (radial) and Y (longitudinal) axes indicate the OCT scanning directions for 2D cross-section and 3D volume measurements, based on left-hand coordinates, with the Z (depth) direction pointing into the page. An electrode used for compound action potential (CAP) recordings was typically placed in the niche near the RWM (see Fig. SI-1 for CAP results). Animal #188R. **b:** A 2D cross-sectional OCT image overlaid with a line drawing of the cells and fluid spaces inferred from the image. The inner boundary of the tectorial membrane (TM) was not entirely visible in the OCT image, so a white dashed outline has been added to show its hypothesized full extent, in which a portion lies just above the IHC stereocilia bundle. (Additional abbreviations found in panels **b–d** are defined further below.) **c:** A labeled drawing of a representative OoC as seen through the RWM, based on the 2D OCT image in **b**. Note that different shades of blue indicate the relative stiffness of different structures, with lighter shades indicating more compliant materials. Cortilymph-filled spaces include the tunnel of Corti (ToC), space of Nuel (SN), outer tunnel (OT), and the spaces between the outer hair cells (OHCs; not shown). The continuation of the BM collagen fibers in the SL region are not entirely visible in this OCT image but are faintly visible in **d**. Note that the dashed lines in SL region have been added to represent these (Agrawal et al., 2018). **d:** A stack of 2D OCT-imaged cross-sections, ranging from ~0.1 to 1.1 mm from the basal end of the cochlea, forms a 3D volumetric representation of the OoC (Movie SI-1 is an animated scan of the image stack). **b-d** Right hand coordinate system. Animal #191. **Other abbreviations:** BCs: Boettcher cells; BM_AZ_ and BM_PZ_: arcuate zone and pectinate zone of the basilar membrane, respectively; CCs: Claudius cells; DCs: Deiters’ cells; HCs: Hensen’s cells; IHC: inner hair cell; IPC and OPC: inner and outer pillar cell, respectively; ISCs: inner-sulcus cells; L: limbus; OSL: osseous (primary) spiral lamina; PhP: phalangeal process; RL: reticular lamina; RM: Reissner’s membrane; SL: spiral ligament; SSL: (osseous) secondary spiral lamina; TBCs: tympanic border cells; TCs: tectal cells; TCE; tectal-cell extension; ST: scala tympani; SM: scala media; SV: scala vestibuli; and ISS: inner spiral sulcus; a.u.: arbitrary units.

While there has been rapid progress in the use of optical coherence tomography (OCT) for in vivo vibrometry measurements of cochlear function, little attention has been paid to the morphology of the OoC in the living cochlea. Nearly all of our knowledge about OoC cellular and extracellular structure comes from histologically processed, postmortem (PM) cochleae that have been decalcified, sliced, stained, and imaged with light or electron microscopy (Lim and Anniko, 1985; O’Malley et al., 2009; Plassmann et al., 1987). Alternate approaches include hemicochlea preparations and confocal microscopy using whole-mount preparations, but they also use PM tissue (Edge et al., 1998; Hardie et al., 2004). Although these are standard methods, their shortcomings include an uncertain relationship to the living anatomy and unknown preparation-related distortions of the fluid spaces and OoC structures. These unknowns have limited our ability to deduce function from anatomy, particularly in the high-frequency basal end of the cochlea that is partially important for sound localization.

The OoC fluid space is defined to contain the following: (1) the inner spiral sulcus (ISS), which is connected to the sub-tectorial fluid space; and (2) the cortilymph space consisting of the tunnel of Corti (ToC), the space of Nuel (SN), and the fluid space around the outer hair cells (OHCs) and outer tunnel (OT; Fig. 1c). In the middle turn of the gerbil cochlea, electrical stimulation of the OHCs has produced fluid waves in the ToC (Karavitaki and Mountain, 2007). Thus, knowledge of the OoC fluid spaces and surrounding structures is important for a full understanding of cochlear amplification (Zagadou et al., 2020). However, these fluid spaces are typically absent in histological images of the high-frequency hook region. As we will show, these absences are likely artifacts due to tissue processing. While many studies have focused on the OoC structures (Hu et al., 1999; Plassmann et al., 1987; Richter et al., 2000; Souter et al., 1995; Spicer et al., 2003), less attention has been paid to the OoC fluid spaces, and particularly the ISS and OT, which brings into question our current understanding of the structure–function relationships in the cochlea at high frequencies. To characterize the in vivo OoC structures and fluid spaces, anatomical methods that work in living animals are needed.

OCT is an optical imaging modality with micron-scale axial and lateral resolutions that enables non-invasive and non-destructive imaging of 2D cross sections and 3D volumes of unlabeled biological tissue using backscattered near-infrared broadband light source (Huang et al., 1991). It is used extensively in a variety of biomedical fields including ophthalmology and otolaryngology, and has additional applications for measuring vibration. OCT has been used to visualize morphological changes in the cochleae of blast- and noise-exposed mice (Kim et al., 2018; Liu et al., 2017), and to visualize changes to the vestibular system in guinea pigs caused by the surgical induction of endolymphatic hydrops (Cho et al., 2015). Micro-OCT (μOCT) technology has been used to image guinea pig temporal bones with sub-micron-level axial and lateral resolutions, allowing visualization of the cochlear microanatomy at a cellular level (Iyer et al., 2016). While the resolutions of μOCT images are the highest yet achieved for OCT imaging, these measurements were from ex vivo cochleae that were chemically fixed.

In this study, we used commercial OCT hardware and custom-built software to collect in vivo 2D images of the gerbil OoC fluid spaces and surrounding structures in the basal-turn hook region. The images were made through the intact round-window membrane (RWM; Fig. 1a, b), with axial (in water) and lateral resolutions of ~1.4 and ~1.95 μm, respectively. In addition, each 3D-volume scans having a depth of about 0.3 mm were obtained at multiple focal depths and then concatenated to obtain about a 1-1.2 mm volume depth (Cho et al., 2018) (Fig. 1d). We obtained in vivo scans over the course of 1 hour, and postmortem scans up to 90 min after death. This procedure allowed us to image the OoC structures while the animal was alive, and to then observe changes in the dimensions and orientations of the OoC structures and fluid spaces during and after the dying process.

The fluid volumes of the ISS, ToC, SN, and OT, as well as the scala tympani (ST) region between the RWM and BM, were segmented for analysis. To directly compare the OCT-based morphometry against morphometry obtained using histological methods, we processed a subset of ears using an aldehyde fixation followed by celloidin embedding and serial sectioning, an approach reported to best preserve the delicate architecture of the inner ear epithelia (O’Malley et al., 2009). We show that many dimensions of the OoC structures, fluid spaces, and cochlear ducts undergo profound changes during the process of death, while others stay constant. Based on these morphological observations, we propose new hypotheses concerning the role of the mechanics of the OT fluid space on OHC transduction, and the effects of the ISS fluid space and sub-tectorial gap on inner hair cell (IHC) and OHC transduction.

## II. Results

We used 3 female and 2 male gerbils for the in vivo studies (N=5), with a subset of these (N=3) also used for the postmortem studies. For each gerbil, in vivo 2D cross-sectional images (e.g., Fig. 1b) were acquired at ~5-min intervals over a 60-min period before the time of death. For three of the gerbils, 2D cross-sectional images continued to be acquired over a 90-min PM period. Using a consistent angular approach, 3D-volume scans (e.g., Fig. 1d, Movie SI-1) were acquired in vivo at the beginning of the experiment and 60 min later, just before death, and at 10-, 30-, 60-, and 90-min after death.

Figure 2 shows representative 2D images at selected time points of the living (top row), and PM (rows 2 and 3) cochlea. The in vivo OoC structures and fluid spaces remained stable across the 60-min interval. In the living cochlea, the RWM and Reissner’s membrane (RM) were curved and bulged outward away from the OoC. A time-lapse movie from the same ear (Movie SI-2) shows major changes occurring in the cochlear anatomy after death. The middle row of Figure 2 shows that both the RWM and RM have started to recede toward the OoC as early as 10-min PM. By 60-min PM, the RWM and RM have changed their directions of curvature, indicating decreased volumes of the ST and scala media (SM). This is consistent with PM decreases in both the ST pressure and the pressure difference between the SM and scala vestibuli (SV). Dimensions of rigid structures such as the outer pillar cells (OPCs) that separate the ToC from the SN (Fig. 1c, Fig. 2 top row) generally remain stable.

**Fig. 2.**
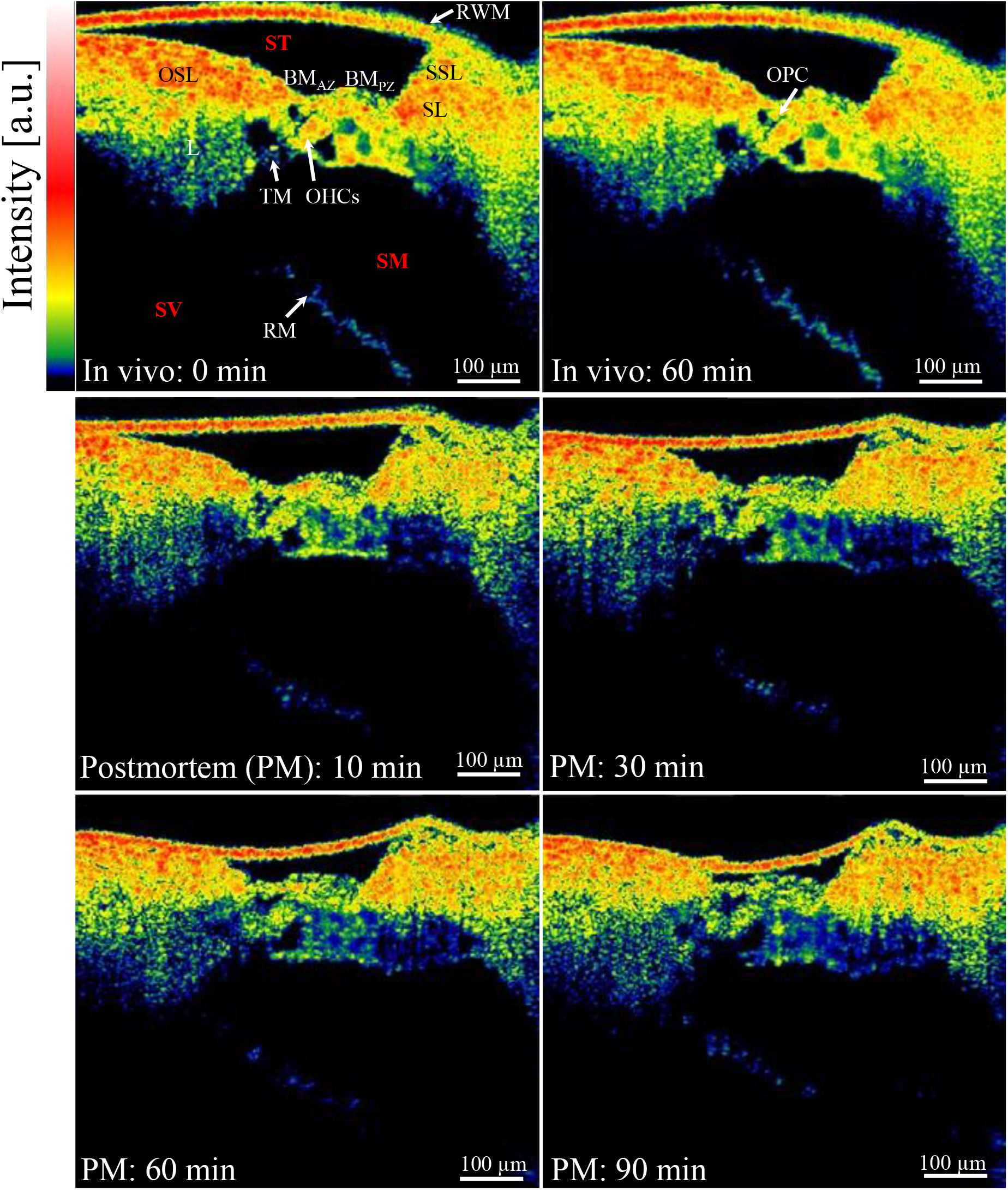
OCT-based cross-sectional OoC images at two in vivo times 60-min apart (top row) and four postmortem (PM) times, at 10, 30, 60, and 90 min after death (rows 2 and 3). The similarities between the in vivo images indicate the stability of the living preparation. The sequence of PM images reveals the progression of changes to the cochlear morphology after death. The full set of cross-sectional images of the PM state, acquired at 5-minute increments, can be viewed as a time-lapse sequence (Movie SI-2). See the caption of Fig. 1 for a list of abbreviations. Animal #269.

Figure 2 qualitatively shows that the OoC structures and fluid spaces changed with death, and that those changes continued to occur up to at least 90-min PM. At 30-min PM, the fluid spaces for the ISS, ToC, SN, and OT appear similar to those at 10-min PM, so the 10-min PM results are not shown in subsequent figures. By 30-min PM, there is a decreased ISS space, an increased OT space, and a swelling of the OoC structure. By 90-min PM the OoC has swelled further (bottom row).

### Quantitative assessment of OoC structural changes from in vivo to postmortem

Figure 3a shows a 2D cross-sectional OCT image illustrating the gross dimensions of the OoC important for cochlear mechanics. To quantify the effects of death on the OoC structure and volume, the OoC width (OoC_W) and height (OoC_H) were measured for the in vivo and PM states and then compared (see SI-3, 4 for methods).

**Fig. 3.**
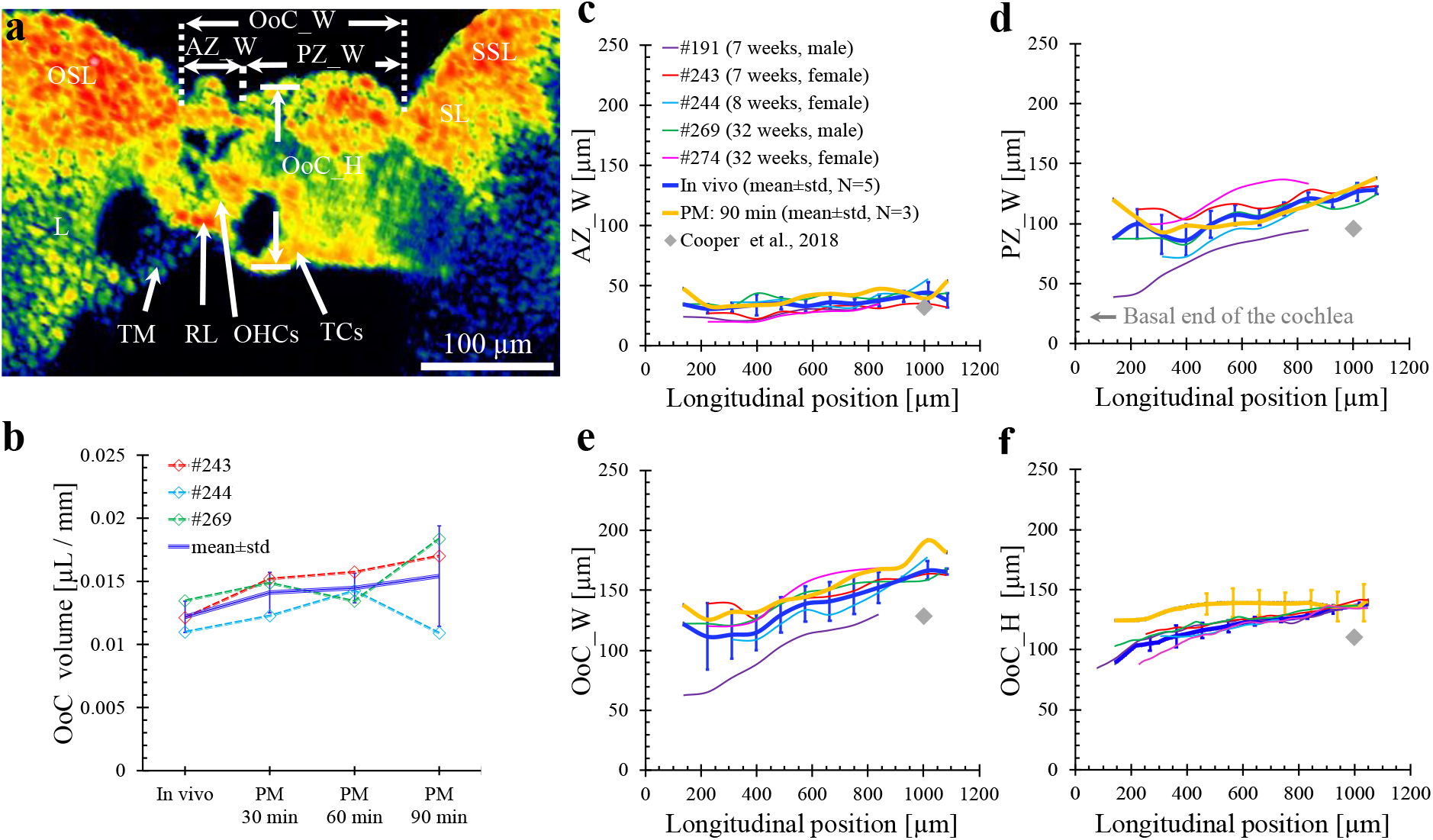
OoC volume and dimensions across in vivo and PM times, and OoC in vivo linear dimensions versus longitudinal position in the cochlear hook region. **a:** This 2D cross-sectional image identifies the BM_AZ_ width (AZ_W) between the OSL ridge and the base of the first row of DCs, the BM_PZ_ width (PZ_W) between the base of the first row of DCs and the SL ridge, and the total OoC width (OoC_W) between the OSL and SL ridges, which is equivalent to the sum of AZ_W and PZ_W. The overall OoC height (OoC_H) is defined as the distance between the BM and the TCs or TCE (see Figs. SI-2 and 3 for segmentation details). Animal #243. **b:** Changes in the OoC volume per unit length (μL/mm) over time, from in vivo to 90-min PM for 3 specimens, along with the mean and standard deviation (std). **c–f:** The AZ_W, PZ_W, OoC_W, and OoC_H dimensions are plotted in respective panels as functions of longitudinal position (in μm) from the basal end of the cochlea. The mean±std error bars for the in vivo measurements (N=5) are shown as blue lines, and those for the 90-min-PM measurements (N=3) are shown as yellow lines. Individual in vivo cases (thin colored lines) are also shown, with the ID, age, and gender of each animal reported in panel **c**. In **c–f**, the gray diamonds are comparison measurements from in vivo OCT images reported by Cooper et al. (2018) for a location about 1 mm (~40 kHz region) from the basal end of the gerbil cochlea.

In the image, OoC_W is bounded by the osseous spiral lamina (OSL) to the left and the spiral ligament (SL) to the right (see also Fig. SI-2). OoC_H is bounded by the BM at the top and a combination of the reticular lamina (RL), tectal cells, and Hensen’s cells at the bottom (see also Fig. SI-2b). The volume of the OoC cellular elements were calculated by subtracting the area of the fluid spaces (see Fig. 5) and integrating the remaining area along the longitudinal dimension. In the region evaluated, the volume per unit length of the structural components of the OoC increased by about 25% by 90-min PM (Fig. 3b). This increase in volume is likely due to swelling of the OoC cells.

The width (AZ_W) and collagen-fiber content of the arcuate zone of the BM (BM_AZ_) are thought to play a dominant role in the place–frequency tonotopic map of the gerbil cochlea (Kapuria et al., 2017). Thus, the total width of the BM (OoC_W) has been divided into AZ_W, bounded by the OSL and the junction of the outer pillar cell and first row base of the Deiters’ cells, and PZ_W, the width of the pectinate zone of the BM (BM_PZ_), which is bounded by the first row base of Deiters’ cells and the SL, as defined previously (see Fig. SI-2a, b, d) (Plassmann et al., 1987; Richter et al., 2000). OoC_W is equivalent to the sum of AZ_W and PZ_W.

Figure 3c–f contains plots of AZ_W, PZ_W, OoC_W, and OoC_H as functions of distance from the basal end of the cochlea toward apex in the hook region over the ~0.1–1.1-mm range. The starting position relative to the basal end of the cochlea was slightly different for each OCT scan (see SI-5 for alignment information). The individual in vivo results are shown (thin colored lines; N=5), along with their mean±standard deviation (std; thick blue lines and error bars), as well as the mean±std (N=3) for the 90-min PM case (thick yellow lines and error bars). OoC_W and OoC_H are slightly larger for the PM state than for the in vivo state. Generally, the dimensions of the OoC increase from the base towards the apex (Fig. 3c–f), which is consistent with the decreasing characteristic frequency in the apical direction that is a key feature of the mammalian cochlea (Peterson and Bogert, 1950; Von Békésy, 1960). Despite the increases in all of these dimensions as one moves apically, the AZ_W/OoC_W ratio maintains an approximately constant value of around 0.25 (not shown). As a comparison, we estimated OoC dimensions based on an in vivo OCT image near the 1-mm location, from Fig. 1c of Cooper et al. (Cooper et al., 2018). These dimensions (gray diamonds in Fig. 3) are typically lower than the averages we found, by about 25%. Our measurements in animal #191 (purple lines), which was the youngest and one of the lower weight animals studied (7 weeks, 72 g), are also lower than our average. The Cooper et al. animal was about 12 weeks and 56 g. Thus, the differences between our results and those of Cooper et al. could be due to lower animal weights, and/or perhaps different OCT-beam angles. The infant and adult and thus age and weight-related OoC dimension differences are known in the human (Meenderink et al., 2019).

### OoC fluid-space changes from in vivo to 90-minutes postmortem

Figure 4 reports the major and minor axes of the OoC fluid subspaces as functions of cochlear location. These subspaces were roughly elliptical in cross section (Fig. 4a). Segmentations of the ISS, ToC, SN, and OT fluid spaces for a representative ear are presented in Figure 4b–e. Figure 4 shows the individual in vivo results (thin colored lines), the corresponding mean±std (N=5, thick blue lines and error bars), and the mean±std for the 90-min PM results (N=3, thick yellow lines and error bars), for the lengths of the major (row i) and minor (row ii) axes. Additionally, the minor/major axis-length ratios are shown for the in vivo (blue lines) and 90-min PM (yellow lines) means (row iii).

**Fig. 4.**
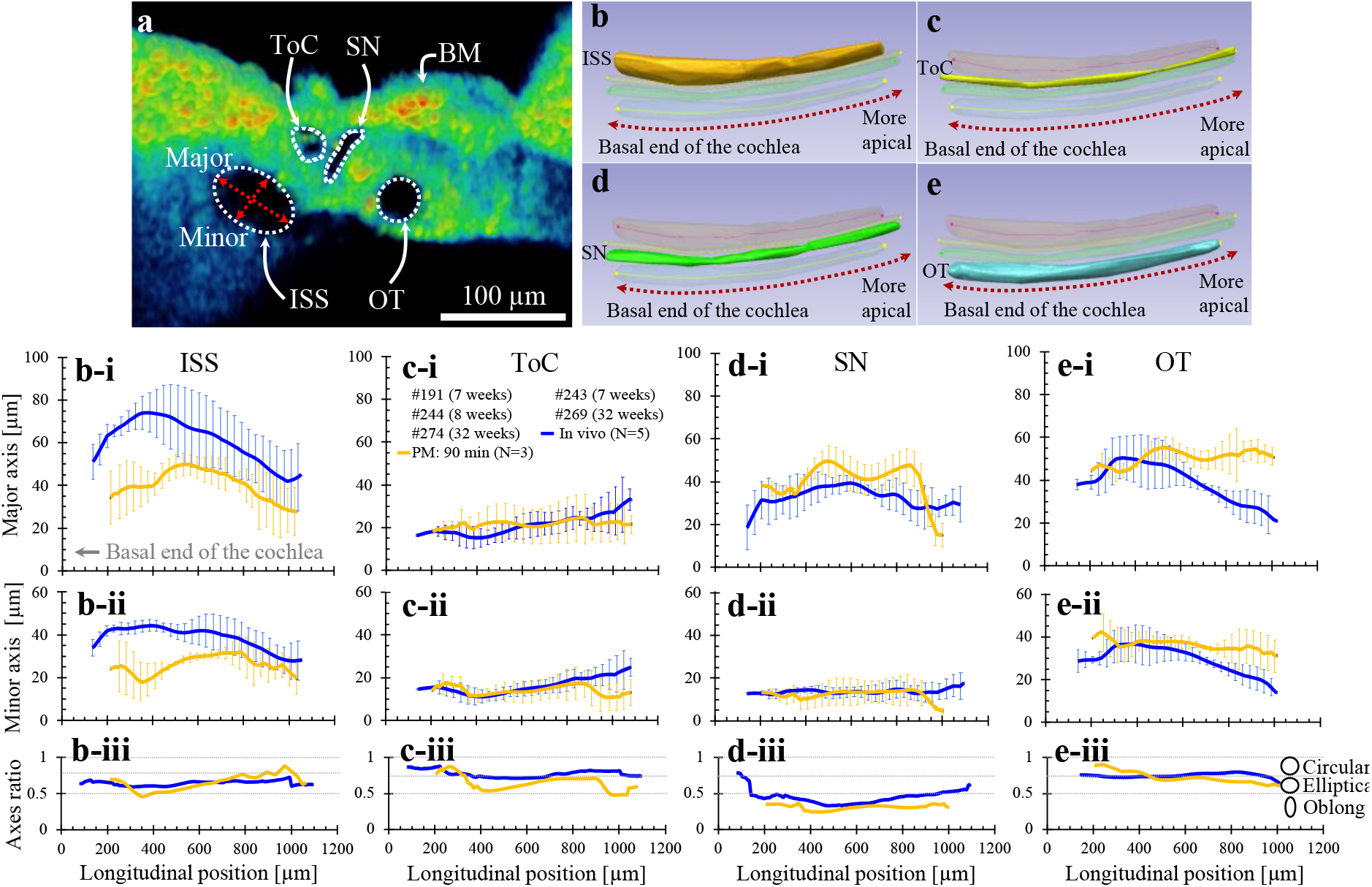
**a:** A 2D OCT cross-sectional image with the BM, ISS, ToC, SN, and OT labeled. Representative in vivo volume segmentations of **b:** inner spiral sulcus (ISS), **c:** tunnel of Corti (ToC), **d:** space of Nuel (SN), and **e:** outer tunnel (OT) are shown for approximately a 1 mm basal section of the cochlea. Animal #191. The inside dimensions of each tubular shape were defined by the major and minor axes of their elliptical cross sections (e.g., red crossed arrows in **a**). Each ellipse was placed orthogonal to the center line that best fit the space at each position (see Fig. SI-4 for segmentation details). **b-i to e-iii:** Inner dimensions of the OoC fluid spaces and minor/major axis-length ratios, for in vivo and 90-min-PM times, as functions of longitudinal position in the cochlear hook region. The mean±std results are in blue for the in vivo cases (N=5) and in yellow for the 90-min-PM cases (N=3). Individual in vivo results are shown as thin colored solid lines, with the ID and age of each animal reported in the legend of panel **c-i**. The second row (**b-i** to **e-i**) shows the corresponding major-axis lengths, while the third row (**b-ii** to **e-ii**) shows the minor-axis lengths. The last row (**b-iii** to **e-iii**) contains the minor/major axis-length ratios.

The ISS fluid space is bounded by the tectorial membrane (TM), limbus (L), and inner-sulcus cells (ISCs), with a narrow opening for the sub-tectorial space (Fig. 1c), generally had the largest in vivo dimensions (Fig. 4).

The average length of the ISS major axis for the in vivo case (Fig. 4b–i) generally increased and then decreased reaching a maximum of about 74 μm. At 90-min PM, the average length of the ISS major axis was smaller. Unlike the major axis, the ISS minor axis for the in vivo case (Fig. 4b–ii) exhibited a flatter longitudinal profile on average, with a value of about 44 μm in the 200–600-μm range. At 90-min PM, the average length of the ISS minor axis was smaller.

In general, the ISS major axis (Fig. 4b-i) showed a similar longitudinal profile between the in vivo and 90-min PM cases, but with the in vivo case scaled higher.

Other OoC dimensions shown in Figure 4 include the ToC (b), SN (c), and OT (d). For the ToC and SN, the PM changes relative to in vivo were minimal. A portion of the OT boundary is formed by tectal cells (Fig. 1b, c), which can be mistaken as Hensen’s cells (Henson et al., 1983). Tectal cells are morphologically distinct from Hensen’s cells, because their membranes form part of the OT fluid space and they are not in contact with the BM (Spicer and Schulte, 1994). The outer wall of the OT is made up of tectal cells, the roof is formed by the tectal-cell extension (TCE), and the medial boundary is formed by the third row of phalangeal processes (PhPs), OHCs, and Deiters’ cells (Fig. 1b, c). The 90-min PM OT dimensions (Fig. 4e-i–ii) generally increased relative to the in vivo case for longitudinal positions apical to the 500-μm location.

The minor/major axis-length ratios indicate the degree to which the cross section of the fluid space is circular (0.75–1.0), elliptical (0.5–0.75), or oblong (<0.5; Fig. 4, bottom row). In general, the ISS (Fig. 4b-iii) was elliptical in cross section and did not change much from in vivo to 90-min PM. The ToC (Fig. 4c-iii) and OT (Fig. 4e-iii) were generally circular for the in vivo case and changed somewhat at 90-min PM. The SN (Fig. 4d-iii) was generally more oblong than the other fluid spaces. Detailed knowledge of the shapes of the OoC fluid spaces will likely be important considerations when constructing finite element (FE) models to better understand the structure-function relationship of the OoC fluid spaces.

To better quantify the progression of PM changes over time, Figure 5 plots the OoC fluid-space volumes for the in vivo state and three PM states 30 min apart. Figure 5a, b shows segmented 3D OCT volume images for a representative animal (#269) for the in vivo (Fig. 5a) and 90-min-PM (Fig. 5b) states. The significant decrease in ST volume (purple) after death is readily apparent. Figure 5c–g contains plots of the ST, ISS, ToC, SN, and OT volumes (normalized by the longitudinal length of the segmented region) as time progresses, for three animals. The volumes of both the ST (Fig. 5c) and ISS (Fig. 5d) decreased systematically from in vivo to 60-min PM. The ISS volume recovered slightly by 90-min PM, whereas the ST volume continued to decrease. The ToC and SN volumes did not change significantly (Fig. 5e, f). On the other hand, the OT volume (Fig. 5g) increased slightly from in vivo to 90-min PM. Table SI-1 lists the corresponding values. PM decreases in the ISS cross-sectional dimensions could be due in part to PM swelling of the sulcus cells. However, this is likely to be a small effect. The decrease in cross-sectional dimensions more likely suggests that the ISS experiences internal pressure in the living cochlea and that this internal pressure decreases during and after death.

**Fig. 5.**
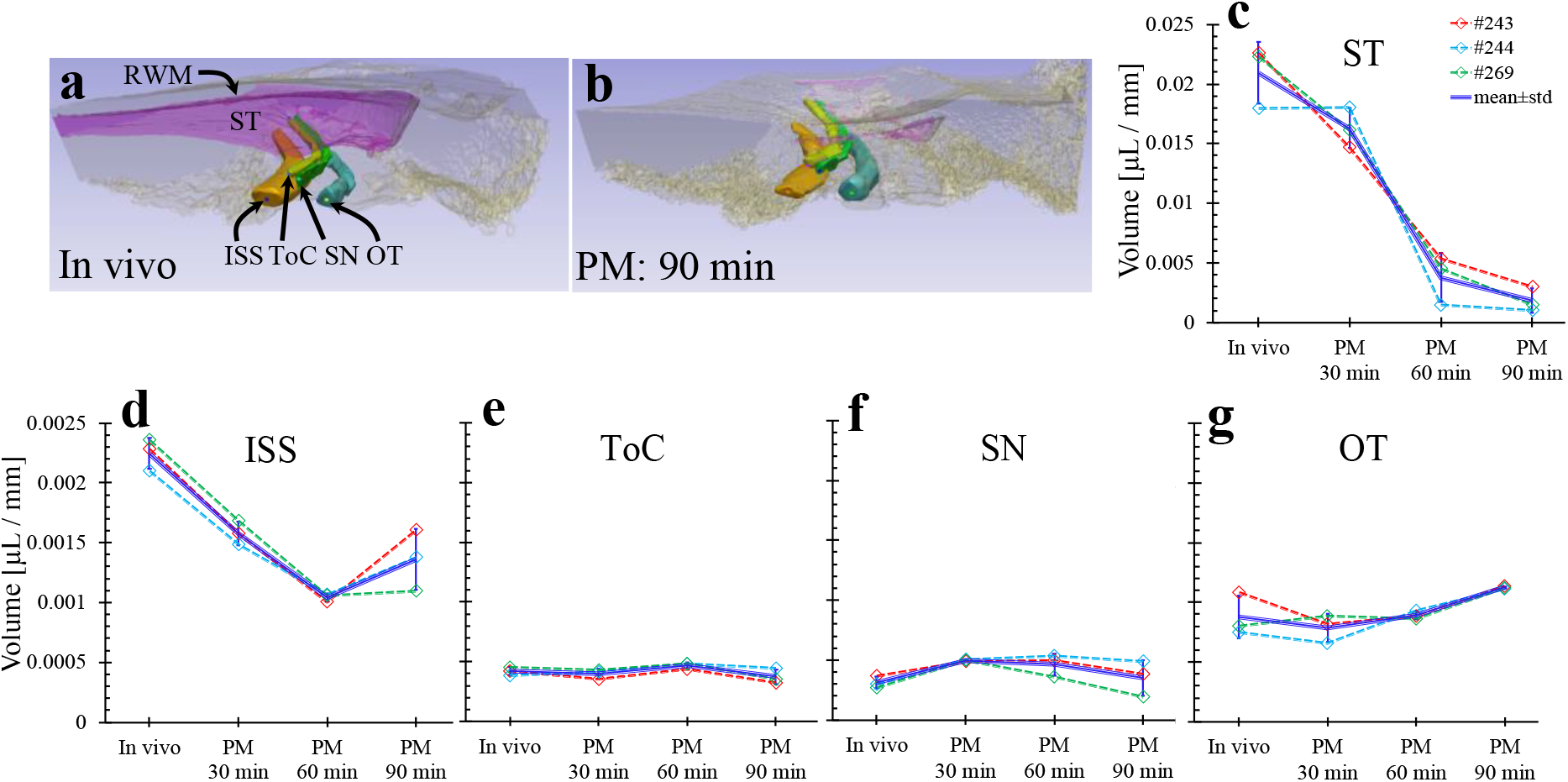
Death-related changes to the ST and OoC fluid volumes. **a–b:** 3D reconstructions of the ST and OoC fluid spaces (ISS, ToC, SN, and OT) based on OCT scans of a representative ear (#269), taken in vivo (**a**), and 90 min after death (**b**). **c–g**: The volume per unit length (μL/mm) is plotted for the in vivo state and each PM time, for the ST (**c**), ISS (**d**; note the change in scale for the second row), ToC (**e**), SN (**f**), and OT (**g**). The mean±std error bars (N=3) are shown as solid-blue lines, and the results from individual ears are shown as colored dashed lines.

### Comparing the structure and fluid-space volumes of the OoC

Figure 6a summarizes the progression of changes in OoC structure and fluid-space volumes from in vivo to 90-min PM. Figure 6a shows that (1) from in vivo to 90-min PM, the average ST volume decreased by more than a factor of 10; (2) the OoC structure alone (without fluid spaces) increased in volume by 25%; and (3) the OoC fluid spaces (all added together) decreased slightly in volume by 20%.

**Fig. 6.**
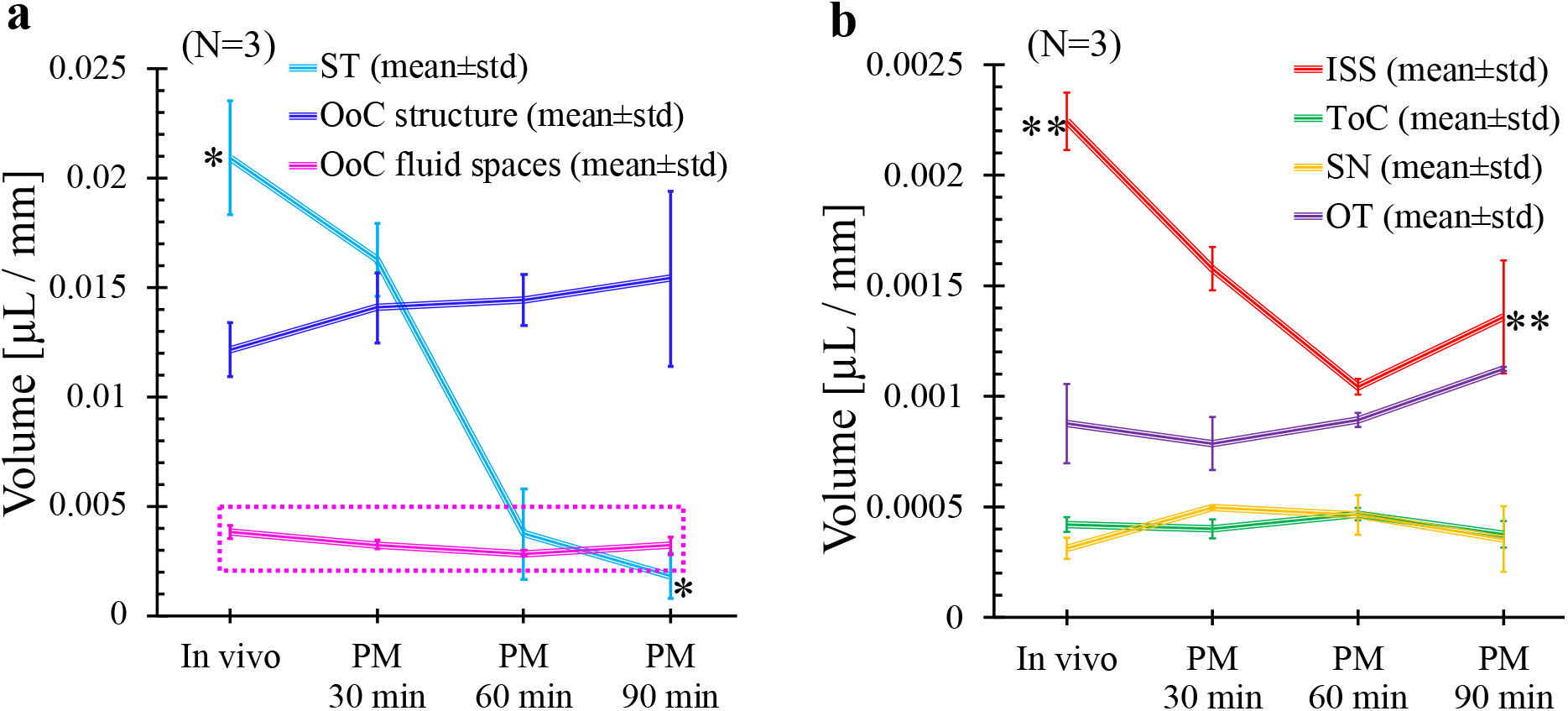
Summary of death-related volume changes in the cochlea. **a:** The mean±std (N=3) of the volumes per unit length (μL/mm) of the ST, the OoC structure (i.e., the OoC volume minus the fluid spaces), and the OoC fluid spaces (i.e., the sum of the ISS, ToC, SN, and OT volumes), plotted for the in vivo and PM times. **b:** The mean±std (N=3) of the volumes per unit length of the individual OoC fluid spaces. The differences between the in vivo and 90-min-PM states were found to be statistically significant for the ST and ISS (\**p* < 0.001 and \*\**p* < 0.01, respectively), as calculated using Student’s t-tests among groups with unequal variances.

Figure 6b summarizes the volume changes of the individual OoC fluid spaces (the ISS, ToC, SN, and OT). Among these, the ISS changed the most after death, decreasing by about 50% from in vivo to 60-min PM (red line). By 90-min PM, the ISS volume had increased slightly. The OT volume (purple line) increased by 20% from in vivo to 90-min PM. Very little change was observed in the smaller ToC (green line) and SN (yellow line) volumes.

The in vivo fluid volumes were statistically compared to each of the PM states using a two-sided Student’s t-test with unequal variances. Only the differences between the in vivo and 90-min-PM volumes of the ST (*; *p* < 0.001) and ISS (**; *p* < 0.01) were statistically significant. The lack of significant changes in the other variables may be due to the small number of animals used.

Some of the PM changes might have resulted from a change in cochlear temperature, which we did not monitor. The animals were placed on a heating pad throughout the measurements, and thus the core body temperature was maintained (See SI-1 for detail).

### OCT vs. histology comparisons

In histological preparations of the cochlea, distortions of the cochlear tissue can arise in multiple ways. In Figures 3–6, we demonstrated and quantified the distortions to the in vivo OoC morphology that take place due to death alone. Additional well-known factors that can distort the OoC structure and fluid spaces include tissue fixation, decalcification, dehydration, embedding, and staining procedures required for histological tissue preparation (Merchant, 2010). To quantitatively assess the degree of combined OoC distortions due to death and histological processing, the OoC morphology was measured after processing with histological methods (N=4).

Figure 7 contrasts two cross-sectional images of the hook region in the same gerbil cochlea (#274): the first was taken in vivo using OCT (Fig. 7a) and the second was obtained through histological methods at approximately the same location (Fig. 7b). The differences between the two images are striking. In Figure 7b (enlarged in Fig. 7c), the TM appears detached, which is sometimes the case with histology, whereas it appears to be normally attached (but less visible) in the in vivo OCT image. In the histological preparation, not only are the SN and ToC fluid spaces hard to distinguish from the OoC structures, but also the OT volume space is dramatically reduced (white dotted circle in Fig. 7c; note the scale bars are different), and the dimensions of the OoC structures appear significantly smaller than those of the in vivo OCT image, especially the OoC_H dimension. In addition, the BMAZ and BMPZ regions are distinguishable in the in vivo OCT image, but a clear boundary between these regions is not visible in the histological image. To verify the dimensions from the histology method, the cochlear volumes from two different animals were segmented, reconstructed, and the outer surfaces aligned (Fig. 7d) based on both a μCT scan (gray) and the histology slices (magenta). The resulting bony volumes overlap, which provides a means of checking that the gross dimensions of the bony structures have not changed and changes observed in the OoC dimensions with histology are due to shrinking of the OoC.

**Fig. 7.**
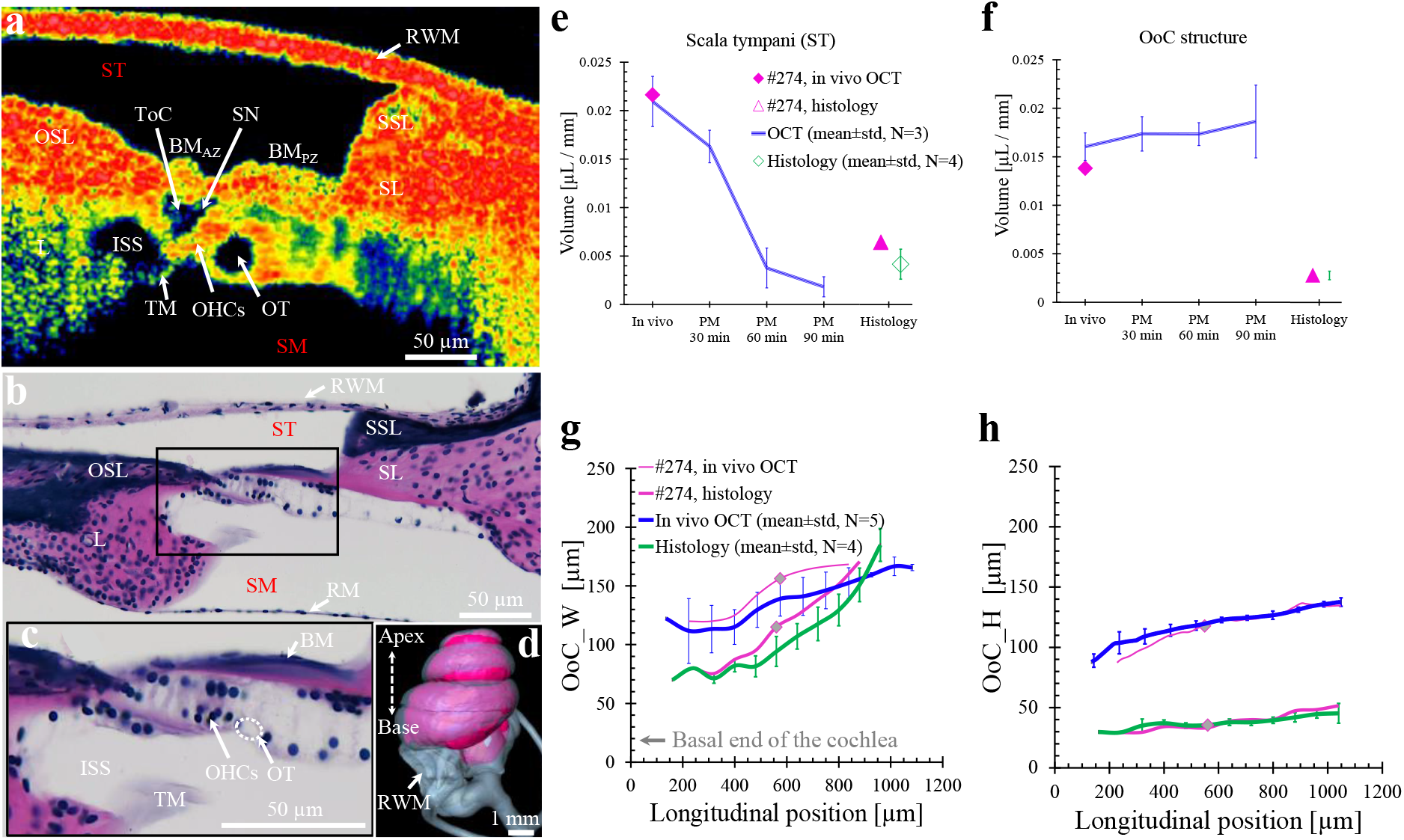
Comparisons of cochlear measurements between OCT (in vivo and PM) and histology (PM only). **a:** An in vivo OCT image shows a cross section of the gerbil OoC ~ 570 μm from basal end of the cochlea. Animal #274. **b:** An image of a histological OoC cross section (#274L-166) from the same ear as in **a**, at a comparable location (about 560 μm from basal end). **c:** An enlarged view of the black box from **b**, showing key OoC structures and the OT (dashed oval; note that the scale bars in **b** and **c** are different). **d:** Cochlear volumes obtained from a μCT scan (gray) and histology (magenta), from two different animals, overlap indicating little change in bony structures due to histology. **e–f:** The volumes per unit length (μL/mm) of the ST fluid space (**e**) and OoC structure (*including* the OoC fluid spaces in this case; **f**) are compared between OCT and histology methods. The OCT results are plotted for the in vivo and PM states in terms of the mean±std (N=3; blue lines), and for the in vivo state of an individual ear (#274; magenta diamonds). The histology results at a comparable location in the same animal (#274, magenta triangles) and as the mean±std (N=4, green diamonds) are plotted to the right. **g–h:** The measured width (OoC_W; **g**) and height (OoC_H; **h**) of the OoC structure are plotted as functions of longitudinal position. The OCT results are plotted for the in vivo mean±std (N=5; blue lines). Note that the gray diamonds correspond to the in vivo OCT and histology measurements of panels **a** and **b-c** respectively. For the OCT method, both the mean±std (blue lines) and individual ear (#274L; thin magenta lines) results are for the in vivo state. The histology measurements are plotted for an individual ear (#274L; thick magenta lines) alongside the mean±std (N=4; green lines).

The OoC dimensions measured with OCT can be compared against those measured by histology. Figure 7e shows the mean±std volume/mm of the ST in multiple animals (N=4) processed with histology (green diamond with error bars), and from an individual ear measured using both OCT and histology (#274; magenta diamond and triangle, respectively). These measurements are compared against the sequence of OCT measurements from in vivo to 90-min PM from Figure 6a (N=3; blue line and error bars). The histology-based ST volume for the individual animal (magenta triangle) was about 50% greater than the mean histology-based volume across ears (green diamond). However, the in vivo OCT-based volume on the same individual ear (magenta diamond) was about three times greater than the histology-based volume. The histology-based volume was comparable to the average of the 60- and 90-min PM volumes (Table SI-2 reports values). Figure 7e shows that a significant amount of the change in ST volume took place between 30- and 60-min PM. For histological processing, the animal is perfused in vivo with a fixative that takes effect within 5 min. In the absence of this perfusion step, however, there was little change in the ST volume during the first 30 min after death (Fig. 7e, blue line). Thus, most of the shrinkage present in the ST volumes in the histological samples likely occurred during this chemical-perfusion stage of histological processing.

Figure 7f compares the volumes (per mm) of the OoC structure (now *including* the OoC fluid spaces) as measured using OCT and histology. The OCT results from in vivo to 90-min PM (N=5; mean±std, blue line) and for one in vivo ear (magenta diamond) are shown alongside the mean±std histology results (N=4, green diamond with error bars concealed by the symbol) and the individual histology results from the same individual in vivo ear (magenta triangle). As mentioned above, the OoC fluid spaces were hard to discern using the histological method, so no attempt was made to subtract those out. The mean volume of the OoC structure (with included fluid spaces) as measured in vivo with OCT (blue line) was about 4.9 times greater than that measured with histology (green diamond), and for the individual ear (magenta diamond vs. triangle) this ratio was about 4.7 times. The volume of the OoC structure was found by integrating the measured cross-sectional areas. These areas were also similarly greater when measured in vivo than when measured with histology (Fig. SI-5a).

Figure 7g and h compare OoC_W and OoC_H, respectively, as measured in vivo vs. in the histological material. Going from the basal end towards the apex, the histology based average OoC_W increases rapidly (Fig. 7g, green line and error bars; N=4). The OCT-based in vivo OoC_W values (Fig. 7g, blue line and error bars; N=5) have a shallower slope than histology and are about 2 times larger than the corresponding histology values at the basal end (near 0.1 mm), but slightly smaller at the more-apical end (near 0.9 mm). Furthermore, the OCT-based in vivo OoC_H values (Fig. 7h) are about 2–3 times larger than those from the corresponding histology measurements. For the individual ear, the differences between the OCT- and histology-based widths (Fig. 7g; thin and thick magenta lines, respectively) appear slightly greater than those between the average widths but also with a steeper slope for histology. The histology-based volume (per mm) and dimensions of the OoC structure are smaller in comparison to the in vivo case (Fig. 7), and the in vivo case is typically smaller than the PM cases (Fig. 3e,f). Recall that for histology, the fixative was in vivo with cardiac perfusion. From this we surmise that the decrease in dimensions in histology (Fig. 7, Fig. SI-5) are not due to PM effects (Fig. 3e,f), but rather due to histological processing. The one caveat is the tangential sectioning angle that in histology could affect the width dimensions, which explains the steeper slope in their width measurements. But the tangential sectioning angle would not affect the height. However, the height with histology decreases quite a bit (Fig. 7g) which again argues that processing decreased the OoC dimensions. The reasons for the smaller width in the basal end are not clear, but this implies that the ridges used to define the OoC_W have moved closer toward each other during histological processing.

## III. Discussion

Our OCT measurements show that anatomical changes take place in the gerbil high-frequency cochlear hook region shortly after death. Some of these changes become apparent within 30-min PM, while others manifest later, e.g., at 90-min PM. Although the advantages of high in-plane resolution as achieved by staining with histology are undeniable, the present OCT-based results demonstrate that our histology-based understanding of the in vivo OoC anatomy has been highly distorted, in the high-frequency hook region. Some of these distortions are due to PM effects, while others are due to histological processing. OCT provides a far more accurate view of in vivo cochlear anatomy and PM effects than has been possible before. The tube-like OT, at the outer border of the RL, and the ISS changed the most from in vivo to 90-min PM. These OCT observations lead us to speculate about the roles of the OT and ISS in OoC mechanics and sound transduction.

A pressure difference across the OoC results in a classic traveling wave and transverse motion of the BM (Fig. 8, red curved vertical arrow), which in gerbils reaches a peak at the BM_AZ_-PZ junction (Cooper et al., 2018). This OoC motion produces pulsatile motion of the cell walls surrounding the OT fluid space (Fig. 8). Depending on the wall stiffness, fluid mass, and steady-streaming Reynolds number, the wall motion may excite resonance modes (Secomb, 1978). We hypothesize that OT resonance modes can induce radial and transverse motions of the TCE, which is attached to the outer edge of the RL and thus provides a mechanism for radial motion of the RL (Fig. 8, red dotted ellipse). Could motion of the OT extracellular fluid space result in a radial RL motion whose phase is important for cochlear amplification? It is generally established that the motion difference between the RL and TM, which causes the OHC stereocilia to deflect, needs to occur with a particular phase relationship to the BM motion so that OHC somatic motility pushes the BM in the direction of BM velocity, such that this motion then opposes viscous losses and results in cochlear amplification (Gummer et al., 1996; Sasmal et al., 2020; Wang et al., 2016). Achieving the correct phase relationship has been attributed to a TM resonance that causes a lag in the radial phase of the TM, starting at about ½-octave below the best frequency (Gummer et al., 1996; Lukashkin et al., 2010; Sasmal and Grosh, 2019). However, recent OCT-based vibrometry measurements have challenged this view, and the TM-resonance hypothesis has been rejected in favor of an alternative hypothesis positing that the source of this special phase relationship is instead a lead in the radial phase of the RL (Guinan, 2020). This hypothesis that the RL provides a leading phase is consistent with the above-best-frequency resonances observed in mouse radial RL and TM motions (Lee et al., 2016). Additional support comes from measurements near the OT in an area called the “lateral compartment”, which show significant in vivo and PM radial motion with the correct phase for triggering OHC somatic motility in the required manner to amplify OoC motion.

**Fig. 8.**
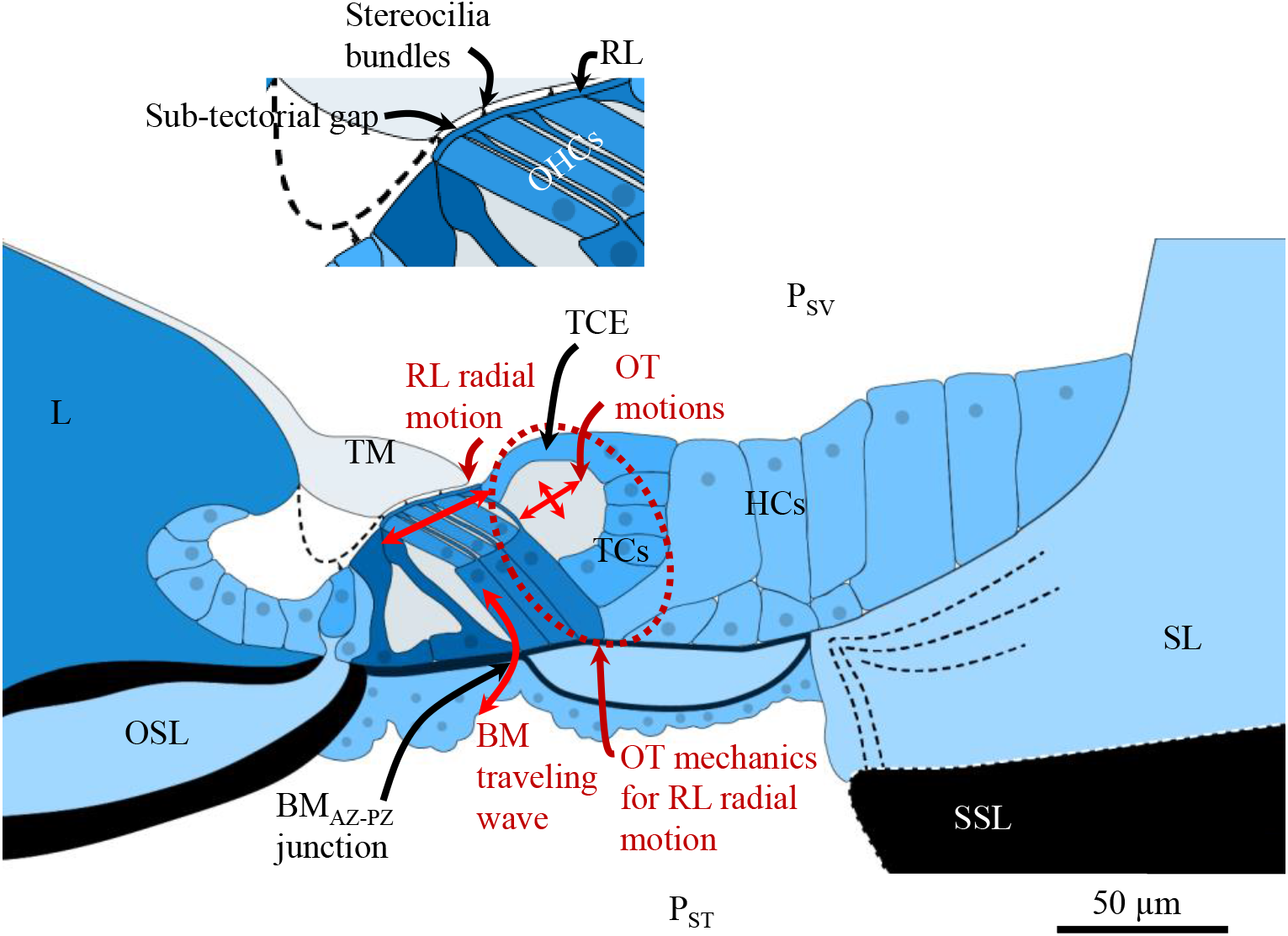
Schematic of the OoC fluid spaces and structures for depicting the proposed OT–RL-resonance hypothesis. The pressure difference between the scala vestibuli (P_SV_) and scala tympani (P_ST_) produces the classic transverse BM traveling wave (red curved vertical arrow at the BM_AZ–PZ_ junction). These cochlear pressures and motions are postulated to affect the OT fluid volume, which may in turn lead to dynamic expansion and contraction (cross-shaped arrows in the OT) of the TCE. The attachment of the TCE to the edge of the RL could then provide a mechanism for radial motion of the RL. Note that the enlarged inset shows details of the sub-tectorial gap and stereocilia bundles between the TM and RL.

However, to our knowledge no mechanism has yet been published that could explain how such a shift in the RL phase could arise. We postulate that the mass of the OT fluid volume, coupled with the stiffness of its surrounding cells, could form an OT–RL resonant structure capable of supporting resonant modes with phase leads in the best-frequency region. This OT–RL resonance would be present for passive mechanics and is hypothesized to be amplified by the active pumping of fluid into the OT by the OHCs.

The ToC, SN, and OT fluid spaces all contain the same cortilymph fluid (gray color, Fig. 8), and have interstitial fluid communication through the spaces around the OHCs and the spaces between the OPCs. It has been observed that electrical excitation of OHC motility causes a pumping of fluid through the SN and into the ToC, and that this possibly enhances cochlear amplification through yet-to-be discovered mechanisms (Karavitaki and Mountain, 2007). A recent FE modeling study features a representation of the gerbil mid-frequency OoC interstitial fluid spaces, and suggests that the role of the interconnected OoC fluid spaces is to redistribute pressure from OHC contractions so as to avoid excessive pressure buildup in the cortilymph space (Zagadou et al., 2020). Comparable FE studies of the high-frequency hook region are currently lacking. Our anatomical measurements will facilitate the building of such models to test the OT– RL-resonance hypothesis presented here.

Does the OT fluid space play a role in longitudinal coupling in the OoC? The OHC-fluid-pumping hypothesis (Karavitaki and Mountain, 2007) predicts the flow of cortilymph through the SN and into the ToC, which would then cause the propagation of a longitudinal wave within the ToC. This fluid wave could interact with the classic BM traveling wave and provide a mechanism for longitudinal coupling between neighboring sections of the OoC (Karavitaki and Mountain, 2007; Zagadou and Mountain, 2012). Thus, using a similar argument as for the ToC, the OT fluid space may also act to promote longitudinal coupling in the OoC. This form of longitudinal coupling would act in addition to other known mechanisms of longitudinal coupling, such as that between neighboring radial collagen fibers of the BM (Von Békésy, 1960), and that caused by the viscoelastic material properties of the TM (Ghaffari et al., 2007; Meaud and Grosh, 2010). Longitudinal coupling in the OoC is also provided by the Y-shaped building blocks that form part of the structure between the RL and BM (each consisting of a Deiters’ cell extending out from the BM, with its PhP and an attached OHC branching out at different angles to join the RL), which are thought to be key for OoC amplification (Motallebzadeh et al., 2018). The relative importance of longitudinal coupling due to wave propagation in the OoC fluid spaces, vs. that due to the motions of structural components of the OoC, has yet to be determined. Such quantitative determinations are likely best approached using computational-modeling efforts.

The OT may also play a role in ion transport, given that the tectal cells of the OT feature fimbriae suggestive of cells active in ion resorption (Spicer and Schulte, 1994). There is K+transport from the base of the OHCs to the Deiters’ cells, and then diffusion via gap junctions between the Deiters’ cells, Hensen’s cells and Claudius cells, back toward the stria vascularis (Kikuchi et al., 2000). It has been proposed that sound-induced K^+^-ion efflux from the OHCs flows into the OT through the interstitial cortilymph fluid (Spicer et al., 2003). These ions would then be absorbed and also channeled through parallel pathways of tectal cells, the third row of Deiters’ cells, and Hensen’s cells, and ultimately find their way toward the stria vascularis (Spicer et al., 2003). The one caveat here is that tectal cells were only seen in the mid to upper turns and not the basal high frequency region (Spicer et al., 2003). However, this result was based on histology studies which distorts the OoC in that region (Fig. 7). Thus, the OT might play a dual role in both OoC mechanics and ion transport.

The decrease in the ISS cross-sectional volume due to the in vivo–PM transition (Figs. 4a, 6b) may have functional significance for IHC and OHC transduction. It is likely that the PM decrease in ISS volume indicates an in vivo regulation of the static pressure within the ISS. Such an in vivo pressure would also maintain a height gap between the TM and RL (Fig. 8, inset image). Because the stiffness of the TM is much less than that of the OHC stereocilia bundles, a sub-tectorial static pressure would be expected to push on the TM to maintain an optimal position just above the top of the tallest row of IHC and OHC stereocilia bundles embedded in the TM. A decrease in ISS static pressure due to death could decrease the sub-tectorial pressure, which would then likely decrease the gap opening and gap height, and thereby possibly disrupt the normal IHC and possibly OHC transduction processes. A change in gap height might also change the angulation of the OHC stereocilia bundles, and thus the bias point for mechano-electric transduction.

Regulation of the sub-tectorial gap opening and gap height due to the pressure in the ISS could be controlled by the aquaporins found in the inner-sulcus cells, which could control the flow of water molecules to and from the ISS (Beitz et al., 2003; Eckhard et al., 2014; Stankovic et al., 1995). Another possible mechanism for pressure regulation is through voltage-regulated ion channels. We postulate that ISS pressurization is necessary to maintain the gap opening near the IHC and the adjoining sub-tectorial gap height between the TM and RL in the living cochlea, and that the loss of this pressurization after death is a contributing factor to the PM loss of cochlear function that has not been previously considered.

From these OCT-based observations, we have postulated a new OT–RL-resonance theory, and a new mechanism by which ISS pressurization regulates the sub-tectorial gap opening and gap height. Testing these hypotheses will require new physiological measurements accompanied by steadily improving OCT methods, as well as further developments in FE modeling approaches.

## IV. Materials and Methods

### Animals and surgical approach

This study was approved by the Institutional Animal Care and Use Committee (IACUC) at Massachusetts Eye and Ear (MEE). Healthy Mongolian gerbils (N=14, aged 5–32 weeks, weight range 60–111 g) of either sex (7 males, 7 females) were used for this study. Results are reported from the five left ears for the OCT and both ears of two animals for histology volume scans were obtained. The surgical procedures and equipment setup are reported in SI-1.

### OCT system and volume scans

All OCT imaging measurements were made using a Spectral Domain–OCT (SD-OCT) system with a 905-nm center wavelength and a 100-kHz high-speed line-scan camera (GAN620C1, Thorlabs, Germany). The system was operated using custom LabVIEW-based software (VibOCT version 1.6). Camera images of the anatomical approach were obtained and used to determine and record the region of interest for the OCT scans (e.g., the black dotted square box in Fig. 1a). VibOCT provides real-time depth-resolved 1D A-scan, 2D cross-sectional B-scan, and 3D volumetric C-scan imaging. The axial resolution was ~1.4 μm (in water, with a refractive index of 1.33), and the lateral resolution was ~1.95 μm, using a 30.5-mm, 0.26-NA long-working-distance, 10× objective lens (M Plan Apo NIR, Mitutoyo, Aurora, IL, USA). To minimize variations among the 3D-volume OCT scans across animals, the orientation of the gerbil head in the head holder, as well as settings on the positioning stage and goniometer, were kept similar across experiments.

While the penetration depth of the OCT system is approximately 1.44 mm (in water), the best B-scan image quality was obtained for depths of less than 300 μm. Because the effective depth of field is smaller than the depth range of interest of about 1-1.2 mm, three or more 3D volume scans were acquired at different focal depths, and tracked by the fine focus adjustment on the OCT optical head with ~1-μm resolution using a Vernier scale (Cho et al., 2018). A single 3D volumetric C-scan consisted of a longitudinal stack of 512 B-scans that are each 1,024 (A-line depth) × 512 (radial), with each C-scan volume averaged 10 times.

All sub-volume scans (each covering a different depth range) were concatenated using ImageJ software to obtain a complete volume stack. The non-overlapping portions from each depth range were extracted, and these portions were then concatenated to obtain a full volume of the OoC in the basal end of the cochlea (the hook region), including the RWM and RM.

All OoC-structure and fluid-space volumes were segmented from the 3D-volume OCT measurements. More details on the 3D-reconstruction and segmentation methods are provided in Figs. SI-2 and 3 for the OoC structure, and in Fig. SI-4 for the OoC fluid spaces.

### Histological preparations

After completing the in vivo 2D and 3D OCT morphological measurements, the anatomies of three gerbils (#192, #274, and #338) were additionally quantified using histological methods, consisting of tissue fixation (using a 4% formaldehyde, 1% acetic acid, 0.1% glutaraldehyde solution) followed by celloidin embedding (O’Malley et al., 2009). Each animal was first intracardially perfused, and then its whole head was extracted, decalcified, and H&E stained, all of which was performed by the Otopathology Laboratory at MEE. Serial sections of the whole skull, including both individual ears, were cut (along the sagittal plane) into 20-μm-thick slices, with every other slice stained to create histology slides. Thus, the histology sections were spaced at 40-μm intervals. The slides were digitized and then reconstructed and segmented into 3D volume stacks for direct comparison against the OoC morphology as obtained using OCT methods (see SI-6 for details).

## Acknowledgements

We thank the MEE Otopathology staff and Anbuselvan Dharmarajan for preparing tissue for histology; Michael E. Ravicz and John J. Guinan, Jr. for technical support and assistance with cochlear-sensitivity measurements; Kevin N. O’Connor, Heidi H. Nakajima, John J. Guinan, Jr., and Rachel Chen for editing assistance; M. Charles Liberman and Jennifer T. O’Malley for helpful discussion with the histology measurements; and Andrew A. Tubelli for the OoC-structure drawings (Fig. 1b, c, and Fig. 8). This work was supported in part by Grant R01 DC07910 from the National Institute on Deafness and Other Communication Disorders (NIDCD) of the NIH, and the Amelia Peabody Charitable Fund.

## Author contributions

N.H.C. and S.P conceived of and designed the project. N.H.C developed the VibOCT software, conducted the experiments, and reconstructed the 3D OCT volumes. H.W performed the segmentation and analysis of the 3D volumes from OCT and histology. S.P, N.H.C, and H.W. wrote and edited the manuscript, and S.P. supervised the project.

## Data availability

All data needed to evaluate the conclusions in the paper are present in the paper. Additional data related to this paper may be requested from the authors.

## Supplementary information

is available for this paper.

## References

Agrawal S, Schart-Morén N, Liu W, Ladak HM, Rask-Andersen H, Li H. 2018. The secondary spiral lamina and its relevance in cochlear implant surgery. Ups J Med Sci 123:9–18. doi:10.1080/03009734.2018.1443983

Beitz E, Zenner H-P, Schultz JE. 2003. Aquaporin-Mediated Fluid Regulation in the Inner Ear. Cell Mol Neurobiol 23:315–329. doi: 10.1023/A:1023636620721

Cho NH, Guan X, Ravicz ME, Puria S. 2018. Human tympanic membrane shape and thickness distribution measured with optical coherence tomography (OCT). Presented at the Asso. Res. Otolaryngology. San Diego, CA, US.

Cho NH, Lee JW, Cho J, Kim J, Jang JH, Jung W. 2015. Evaluation of the usefulness of three-dimensional optical coherence tomography in a guinea pig model of endolymphatic hydrops induced by surgical obliteration of the endolymphatic duct. Journal of biomedical optics 20:036009.

Cooper NP, Vavakou A, van der Heijden M. 2018. Vibration hotspots reveal longitudinal funneling of sound-evoked motion in the mammalian cochlea. Nat Commun 9:3054. doi:10.1038/s41467-018-05483-z

Eckhard A, Müller M, Salt A, Smolders J, Rask-Andersen H, Löwenheim H. 2014. Water permeability of the mammalian cochlea: functional features of an aquaporin-facilitated water shunt at the perilymph–endolymph barrier. Pflugers Arch - Eur J Physiol 466:1963–1985. doi:10.1007/s00424-013-1421-y

Edge RM, Evans BN, Pearce M, Richter C-P, Hu X, Dallos P. 1998. Morphology of the unfixed cochlea. Hearing research 124:1–16.

Ghaffari R, Aranyosi AJ, Freeman DM. 2007. Longitudinally propagating traveling waves of the mammalian tectorial membrane. Proc Natl Acad Sci U S A 104:16510–16515. doi:10.1073/pnas.0703665104

Guinan JJ. 2020. The interplay of organ-of-Corti vibrational modes, not tectorial-membrane resonance, sets outer-hair-cell stereocilia phase to produce cochlear amplification. Hearing Research 395:108040. doi:10.1016/j.heares.2020.108040

Gummer AW, Hemmert W, Zenner HP. 1996. Resonant tectorial membrane motion in the inner ear: its crucial role in frequency tuning. Proceedings of the National Academy of Sciences 93:8727–8732. doi:10.1073/pnas.93.16.8727

Hardie NA, MacDonald G, Rubel EW. 2004. A new method for imaging and 3D reconstruction of mammalian cochlea by fluorescent confocal microscopy. Brain Research 1000:200–210. doi:10.1016/j.brainres.2003.10.071

Henson MM, Jenkins DB, Henson OW. 1983. Sustentacular cells of the organ of Corti — the tectal cells of the outer tunnel. Hearing Research 10:153–166. doi:10.1016/0378-5955(83)90051-5

Hu X, Evans BN, Dallos P. 1999. Direct visualization of organ of Corti kinematics in a hemicochlea. Journal of neurophysiology 82:2798–2807.

Huang D, Swanson EA, Lin CP, Schuman JS, Stinson WG, Chang W, Hee MR, Flotte T, Gregory K, Puliafito CA. 1991. Optical coherence tomography. science 254:1178–1181.

Iyer JS, Batts SA, Chu KK, Sahin MI, Leung HM, Tearney GJ, Stankovic KM. 2016. Micro-optical coherence tomography of the mammalian cochlea. Sci Rep 6:33288. doi: 10.1038/srep33288

Kapuria S, Steele CR, Puria S. 2017. Unraveling the mystery of hearing in gerbil and other rodents with an arch-beam model of the basilar membrane. Scientific Reports 7:228. doi: 10.1038/s41598-017-00114-x

Karavitaki KD, Mountain DC. 2007. Evidence for outer hair cell driven oscillatory fluid flow in the tunnel of Corti. Biophysical journal 92:3284–3293.

Kikuchi T, Adams JC, Miyabe Y, So E, Kobayashi T. 2000. Potassium ion recycling pathway via gap junction systems in the mammalian cochlea and its interruption in hereditary nonsyndromic deafness. Medical Electron Microscopy 33:51–56. doi:10.1007/s007950070001

Kim J, Xia A, Grillet N, Applegate BE, Oghalai JS. 2018. Osmotic stabilization prevents cochlear synaptopathy after blast trauma. Proceedings of the National Academy of Sciences 115:E4853–E4860.

Lee HY, Raphael PD, Xia A, Kim J, Grillet N, Applegate BE, Ellerbee Bowden AK, Oghalai JS. 2016. Two-Dimensional Cochlear Micromechanics Measured In Vivo Demonstrate Radial Tuning within the Mouse Organ of Corti. Journal of Neuroscience 36:8160–8173. doi:10.1523/JNEUROSCI.1157-16.2016

Lim DJ, Anniko M. 1985. Developmental Morphology of the Mouse Inner Ear: A scanning electron microscopic observation. Acta Oto-Laryngologica 99:5–69. doi:10.3109/00016488509121766

Liu GS, Kim J, Applegate BE, Oghalai JS. 2017. Computer-aided detection and quantification of endolymphatic hydrops within the mouse cochlea in vivo using optical coherence tomography. Journal of Biomedical Optics 22:076002.

Lukashkin AN, Richardson GP, Russell IJ. 2010. Multiple roles for the tectorial membrane in the active cochlea. Hearing Research 266:26–35. doi:10.1016/j.heares.2009.10.005

Meaud J, Grosh K. 2010. The effect of tectorial membrane and basilar membrane longitudinal coupling in cochlear mechanics. J Acoust Soc Am 127:1411–1421. doi:10.1121/1.3290995

Meenderink SWF, Shera CA, Valero MD, Liberman MC, Abdala C. 2019. Morphological Immaturity of the Neonatal Organ of Corti and Associated Structures in Humans. JARO 20:461–474. doi:10.1007/s10162-019-00734-2

Merchant SN. 2010. Schuknecht’s Pathology of the Ear. PMPH-USA.

Motallebzadeh H, Soons JAM, Puria S. 2018. Cochlear amplification and tuning depend on the cellular arrangement within the organ of Corti. Proc Natl Acad Sci USA 115:5762–5767. doi:10.1073/pnas.1720979115

O’Malley JT, Merchant SN, Burgess BJ, Jones DD, Adams JC. 2009. Effects of Fixative and Embedding Medium on Morphology and Immunostaining of the Cochlea. Audiol Neurootol 14:78–87. doi:10.1159/000158536

Peterson LC, Bogert BP. 1950. Erratum: A Dynamical Theory of the Cochlea. The Journal of the Acoustical Society of America 22:640–640. doi:10.1121/1.1906668

Plassmann W, Peetz W, Schmidt M. 1987. The Cochlea in Gerbilline Rodents. BBE 30:82–102. doi:10.1159/000118639

Richter C-P, Edge R, He DZZ, Dallos P. 2000. Development of the Gerbil Inner Ear Observed in the Hemicochlea. JARO 1:195–210. doi:10.1007/s101620010019

Sasmal A, Geib N, Popa B-I, Grosh K. 2020. Broadband nonreciprocal linear acoustics through a non-local active metamaterial. New J Phys 22:063010. doi:10.1088/1367-2630/ab8aad

Sasmal A, Grosh K. 2019. Unified cochlear model for low- and high-frequency mammalian hearing. Proc Natl Acad Sci USA 116:13983–13988. doi:10.1073/pnas.1900695116

Secomb TW. 1978. Flow in a channel with pulsating walls. Journal of Fluid Mechanics 88:273–288. doi:10.1017/S0022112078002104

Souter M, Nevill G, Forge A. 1995. Postnatal development of membrane specialisations of gerbil outer hair cells. Hearing research 91:43–62.

Spicer SS, Schulte BA. 1994. Ultrastructural differentiation of the first hensen cell in the gerbil cochlea as a distinct cell type. The Anatomical Record 240:149–156. doi:https://doi.org/10.1002/ar.1092400202

Spicer SS, Smythe N, Schulte BA. 2003. Ultrastructure indicative of ion transport in tectal, Deiters, and tunnel cells: Differences between gerbil and chinchilla basal and apical cochlea. Anat Rec 271A:342–359. doi:10.1002/ar.a.10041

Stankovic KM, Adams JC, Brown D. 1995. Immunolocalization of aquaporin CHIP in the guinea pig inner ear. American Journal of Physiology-Cell Physiology 269:C1450–C1456. doi:10.1152/ajpcell.1995.269.6.C1450

Von Békésy G. 1960. Experiments in hearing., McGraw-Hill series in psychology. New York: McGraw.

Wang Y, Steele CR, Puria S. 2016. Cochlear Outer-Hair-Cell Power Generation and Viscous Fluid Loss. Sci Rep 6:19475. doi:10.1038/srep19475

Zagadou BF, Barbone PE, Mountain DC. 2020. Significance of the Microfluidic Flow Inside the Organ of Corti. Journal of Biomechanical Engineering 142:081009. doi:10.1115/1.4046637

Zagadou BF, Mountain DC. 2012. Analysis of the Cochlear Amplifier Fluid Pump Hypothesis. JARO 13:185–197. doi:10.1007/s10162-011-0308-x

