## Supplemental Information for "Watching Death in the Gerbil Cochlea Using Optical Coherence Tomography"

**File name**: “Movie SI-1.mov”. Description: A slice-by-slice traversal of the OCT-based 3D reconstruction of an in vivo gerbil organ of Corti (OoC) in the hook region, as described in Fig. 1d.

**File name**: “Movie SI-2.mov”. Description: A time-lapse view of the effects of death on a cross section of the OoC in the hook region, as described in Fig. 2.

### 1. Surgical approach and equipment setup

All surgical procedures, including anesthesia, were performed in a private surgery room near the sound-proof chamber in which the experiments were conducted. The initial anesthesia was induced by an intraperitoneal (IP) injection of sodium pentobarbital (70 mg/kg), followed by a subcutaneous (SQ) injection of acepromazine (1 mg/kg) mixed with atropine (0.06 mg/kg). Lidocaine (1%) with epinephrine (1:100,000) was applied topically to the skin over the top of the skull and around the left ear. Puralube was applied to the eyes to reduce the drying effects of the microscope light during surgery. An SQ or IP injection of Ringer's solution (1 mL/kg) was performed every 1 hour to maintain fluid volume, and as necessary to replace lost fluid (for example, due to bleeding). To maintain adequate anesthesia, 1/3 of the initial dose of sodium pentobarbital was given approximately every 45–60 minutes, and the depth of anesthesia was evaluated every 30–60 minutes via toe-pinch response and/or an increase of more than 10% in the monitored heart rate. Half of the acepromazine initial dose was injected approximately every 3–4 hours. Atropine (the full initial dose) was reinjected after 2–3 hours in case of respiratory difficulty.

The gerbils were generally able to self-ventilate, but if an animal appeared to have difficulty breathing, a tracheostomy was performed, and a ventilation tube was inserted into the trachea to ensure breathing without obstruction. Once the gerbil was under deep anesthesia, it was placed on a thermostatically controlled heating pad to maintain a rectal temperature of 38.5±1 ºC. The head of the gerbil was firmly fixed using a customized head holder. The pinna and external ear canal of the left ear were removed, and the tympanic bulla on the same side was exposed through a ventral approach in the same surgical field used for the tracheostomy, with care taken throughout to minimize noise and vibration in order to avoid mechanically induced hearing damage.

The diameter of the opening in the posterolateral wall of the bulla was gradually increased using a small (#15) sharp knife. The tympanic membrane, malleus, incus, stapes, and round-window membrane (RWM) were all kept intact. Subcutaneous electrodes were placed for continuous electrocardiogram (EKG) monitoring. A silver-ball electrode was placed in the round-window niche, and a ground electrode was placed in the soft tissue of the neck or attached to the head-holder, in order to record the compound action potential (CAP).

After completing the surgical procedures, the gerbil was carefully transferred to the sound-proof chamber to begin OCT imaging. The animal, heating pad, EKG electrode, and surgical bed were transferred to a two-stage goniometer (07-GON-503, Melles-Griot, Carlsbad, CA, USA), which was positioned on top of a 3-axis micro-manipulator (OCT-XYR1, Thorlabs, Germany). All of these were mounted on a vibration-isolation table. The in vivo and PM imaging measurements were performed in the hook region of the cochlea through the intact RWM.

After completing the in vivo imaging and physiological measurements, an IP injection of Fatal Plus (>150 mg/kg) was administered for euthanasia. The heart typically stops within 5–10 minutes after injection, at which point the animal stops breathing. The reported PM time points are in relation to the moment when the animal stopped breathing, had no heartbeat, and exhibited no other movements.

### 2. Monitoring cochlear sensitivity

The physiological condition of the cochlea was monitored in vivo using CAP responses to a series of clicks at different sound levels (20–80 dB SPL, in 10-dB steps), collected both before and after the in vivo 3D volume scan. The acoustic clicks were produced by a speaker (Beyerdynamic DT-48; Heilbronn, Germany) and coupled to the ear canal ring. A calibrated-microphone reading (Knowles FG-23629; Itasca, IL, USA) of the sound pressure in the ear canal was used during each experiment to confirm the sound levels. Signal generation and sound-pressure control for the CAP recordings were accomplished using an NI PXI-4461 dynamic signal acquisition device mounted in an NI PXI-1031 chassis with an NI PXI-8196 embedded computer (National Instruments; Austin, TX, USA). The hardware was controlled using the custom LabVIEW-based Cochlear Function Test Suite (EPL_CFTS, version 2333, (Katsumi et al., 2020)). The CAP signal was amplified by a P511 amplifier (Grass Instruments; Warwick, RI, USA), and $|N_1–P_1|$ was calculated in the software.

Figure SI-1 illustrates cochlear-sensitivity monitoring using click-evoked CAP readings. Figure SI-1a demonstrates very robust and clean CAP readings (#274), with $N_1$ to $P_1$ growing over a large dynamic range (30–70 dB SPL, shown in 10-dB steps). The click SPL corresponding to a CAP reading with a peak-to-peak $|N_1–P_1|$ amplitude of 15 µV was defined as the "threshold" CAP response in this study. The average threshold values across all four gerbils were 44.5 and 41 dB SPL, respectively, based on readings taken before and after the in vivo 3D-volume measurements. These values indicate that the health of the cochlea was maintained throughout the in vivo state (Fig. SI-1b).

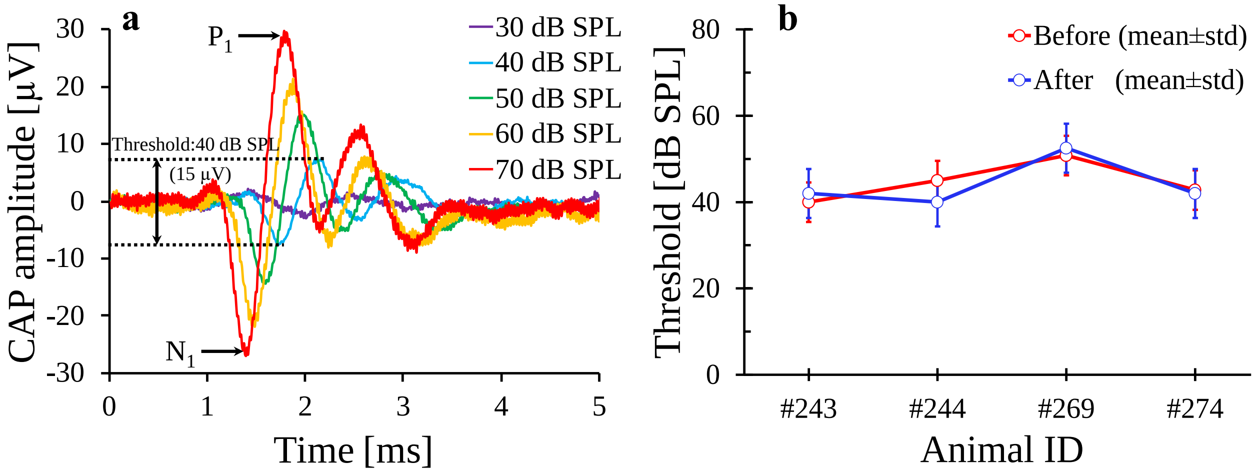


**Fig. SI-1.** Cochlear sensitivity as monitored using sound-evoked compound action potentials (CAPs). **a:** The CAP responses from 30 to 70 dB SPL clicks, plotted as functions of time for one gerbil (#274). The "threshold" was defined as the click SPL that came closest to evoking a 15-µV peak-to-peak CAP (black dotted lines) from the first negative peak ($N_1$) to the first positive peak ($P_1$), i.e., $|N_1 – P_1|$. **b:** The threshold means and standard deviations (std) for CAP responses from four in vivo gerbils, before the first 3D OCT volume measurements (red line) and at the end of the experiment before euthanasia (blue line). Thresholds were measured several times on 4 of the 5 animals, but CAP measurements were not made on the first animal (#191).

### 3. Segmenting the organ of Corti (OoC)

The software applications Amira (version 2019.1; Thermo Fisher Scientific; Waltham, MA, USA) and SimpleWare ScanIP (version P-2019.09; Synopsis; Exeter, UK) were used to acquire 3D reconstructions and segmentations of the cochlear structures, respectively. The images in the OCT volume stacks were segmented semi-automatically using a combination of thresholding, morphological operations (i.e., dilation, erosion, etc.), as well as smoothing and interpolation tools, then reconstructed into 3D objects in both SimpleWare and Amira (more detail below in Section 4). The segmented objects were then analyzed to extract the total width and height of the OoC (OoC_W and OoC_H, respectively), the respective widths of the arcuate (AZ_W) and pectinate

(PZ_W) zones of the BM, and the shapes and volumes of the following OoC fluid spaces: the inner spiral sulcus (ISS), tunnel of Corti (ToC), space of Nuel (SN), and outer tunnel (OT).

Figure SI-2c shows a reconstructed 3D representation of a portion of the hook region that extends about 1.1 mm from the basal end of the cochlea (white dotted arrow), assembled from stacks of 1,024 × 512 2D OCT cross sections, such as the one shown in Fig. SI-2b. The OoC structure is shown as the orange region between the osseous spiral lamina (OSL) and spiral ligament (SL) ridge. Figure SI-2d shows a top-down view of the segmented OoC structure, with the RWM and secondary spiral lamina (SSL) hidden and a white dividing line added between the arcuate-zone ($\mathrm{BM_{AZ}}$) and pectinate-zone ($\mathrm{BM_{PZ}}$) regions of the BM. The boundary between the $\mathrm{BM_{AZ}}$ and $\mathrm{BM_{PZ}}$ regions coincides with the location of the junction of the outer pillar cell (OPC) and first row base of the Deiters' cells (1$^{st}$ row of DC; Fig. SI-2a and b). OoC_W is the sum of AZ_W and PZ_W (Fig. SI-2a and b).

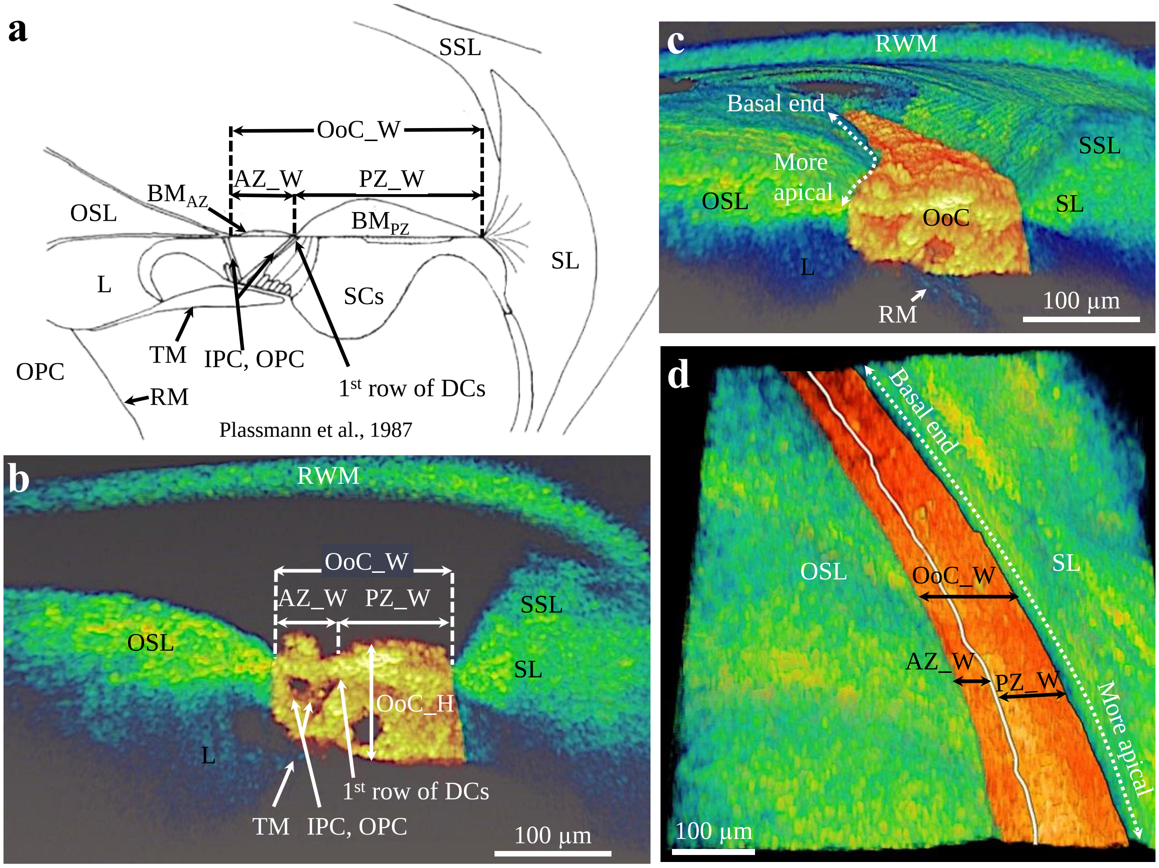


**Fig. SI-2.** Segmenting the organ of Corti (OoC) in the hook region and determining OoC_W, AZ_W, and PZ_W. **a:** A cross-sectional drawing of the OoC (adapted from (Plassmann et al., 1987)), showing the definitions of the widths of the OoC (OoC_W), the arcuate (AZ_W) and pectinate (PZ_W) zones of the basilar membrane (BM); and identifying the osseous (primary) spiral lamina (OSL), limbus (L), Reissner's membrane (RM), tectorial membrane (TM), inner and outer pillar cells (IPC, OPC), the first row of Deiters' cells (DCs), support cells (SCs), and spiral ligament (SL). **b:** An in vivo 2D OCT cross-sectional image showing the OoC structure (orange color), with the defined widths (OoC_W, AZ_W, and PZ_W) and height (OoC_H) labeled, as well as the RWM and some of the other structures shown in **a**. **c:** A 3D reconstruction based on in vivo OCT volume scans, indicating the segmented OoC structure in the hook region (orange color) between the OSL and SL ridge, from the basal end toward a more-apical position (white dotted line). **d:** A view of the segmented OoC structure with a white line dividing the AZ_W and PZ_W subdivisions of the BM, and with the RWM and (osseous) secondary spiral lamina (SSL) removed for better visualization of the segmented OoC structure. **b-d:** Animal #191.

### 4. Determining OoC_W and OoC_H from OCT Data

The total width of the OoC as reported in Fig. 3e, OoC_W, was determined by computing the sum of AZ_W (Fig. 3c) and PZ_W (Fig. 3d). These measurements were conducted within the OCT-imaging planes. For comparison, we also determined OoC_W within planes orthogonal to the BM centerline.

Figure SI-3a shows a 2D cross-sectional OCT image with a thin BM layer (red region with white dotted outline) segmented using the volume-editing tool in Amira. Figure SI-3b shows a 3D view of the thin BM layer (in red), from the basal end to a more-apical location, with the rest of the segmented OoC structure shown in gray. Figure SI-3c shows the BM layer by itself. The segmented BM was then imported into SimpleWare to calculate the centerline and fit circumscribed circles perpendicular to the centerline. Figure SI-3d shows an example circumscribed circle fit to the BM layer. The diameters of these circles, spaced at 5-voxel (8 µm) intervals (Fig. SI-3e), represent the version of OoC_W that is discussed in SI-9.

The height of the OoC as reported in Figure 3f, OoC_H, was solely determined using a inscribed circle fitting procedure. The segmented OoC structure (Fig. SI-2c) was imported into SimpleWare to calculate the centerline and fit inscribed circles perpendicular to the centerline. Figure SI-3f shows an example inscribed circle. The diameters of these inscribed circles, also spaced at 8-µm intervals (Fig. SI-3g) represent OoC_H.

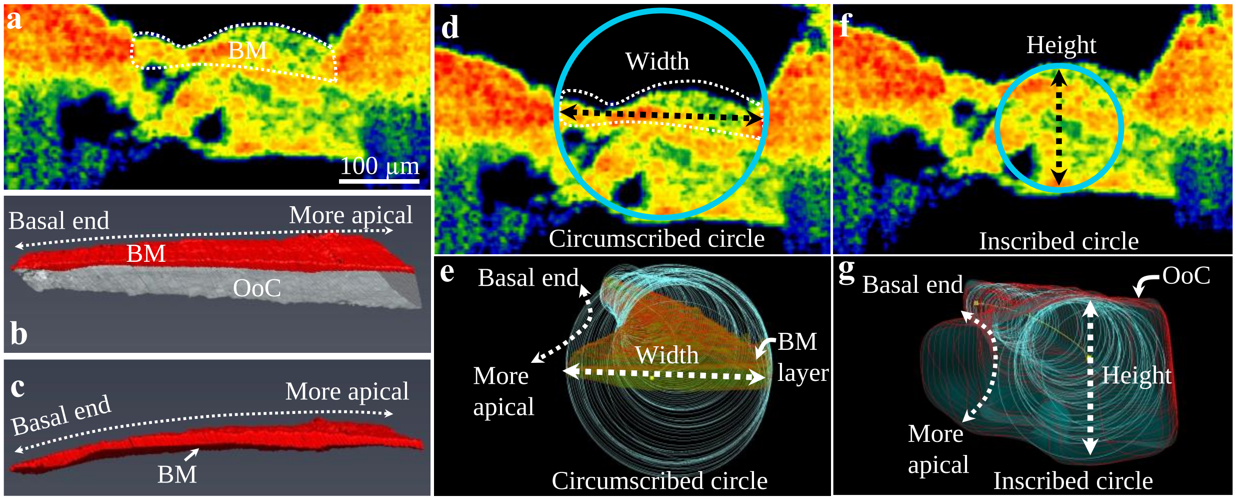


**Fig. SI-3.** Determining the width and height of the OoC along the cochlear length by fitting circles along the centerline in 3D. **a:** A 2D OCT cross-sectional image of the OoC with the BM layer inside the white-dotted outline. **b:** The segmented 3D volume of the OoC structure (gray) is shown with the segmented BM volume layer (red) on top. **c:** The extracted thin BM-volume layer. **d:** The thin BM layer (white-dotted outline) was fit with circumscribed circles (light blue) perpendicular to the centerline, the diameters of which provide an alternate way of determining OoC_W using a circle fitting method that is not tied to the OCT imaging planes. **e:** The segmented 3D volume of the BM layer showing the circumscribed circles (light blue; spaced 8-µm apart along the centerline) along the length of the OoC that represent OoC_W. **f:** The segmented OoC structure was fit with inscribed circles (light blue) perpendicular to the centerline, the diameters of which determine OoC_H. **g:** The segmented 3D volume of the OoC is shown with the inscribed circles (light blue; spaced 8-µm apart along the centerline) along the length of the OoC that represent representing OoC_H. **a-g:** Animal #269.

### 5. Segmenting the OoC fluid spaces and measuring their inner dimensions

The starting position relative to the basal end of the cochlea was slightly different for each OCT scan. To estimate the starting position for each individual ear, we used as a reference a whole-cochlea 3D reconstruction from a µCT scan of a different specimen. In Amira, the centerline of the OoC of this whole-cochlea reconstruction was then used to align and register the segmented OCT volumes at the most appropriate location in the hook region by visually aligning in 3D using the round-window opening as a landmark and the curvature of the OoC structure for guidance. Figure SI-4a shows an example of an OCT-based segmented OoC-structure volume (the red region within the white dotted ellipse) whose centerline has been registered with the whole-cochlea OoC centerline. By aligning the centerlines, the estimated starting points of the OCT-based volumes ranged from 78 to 228 μm from the basal end of the cochlea, with each volume extending approximately 1.1 mm in the apical direction. Centerlines were also determined for the fluid-space volumes, such that circumscribed ellipses orthogonal to the centerline were fit to the shape of each cross section along the length of the volume.

The major and minor axes of the ellipses were then used to gauge the varying cross-sectional shapes of each fluid space. Figure SI-4b shows the OCT-based 3D reconstruction from the basal end, including segmented representations of the ST, ISS, ToC, SN, and OT fluid spaces. Figure SI-4c shows an OCT cross-section of the OoC with the fluid spaces indicated. Figure SI-4d shows the segmented OoC fluid spaces on their own, each with a visible centerline used for fitting ellipses to each volume along its length in order to gauge the changing shape of the volume in terms the major and minor axes of the ellipses (e.g., the red arrows on the ISS). The centerlines were calculated in SimpleWare based on the Skeletonization algorithm. Figure SI-4e provides an alternate view of the OoC fluid spaces.

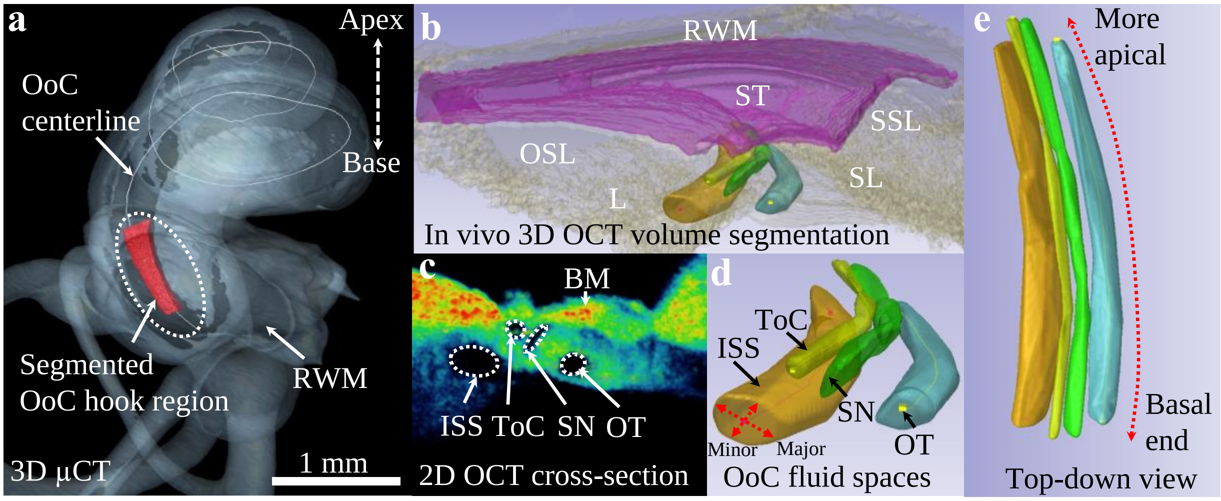


**Fig. SI-4.** Segmentation of the OoC fluid spaces. **a:** The OCT-based segmented region of the OoC (red) is shown superimposed upon the hook region of a full μCT-based reconstruction of the cochlea (gray). **b:** The scala tympani (ST) and OoC fluid spaces (from the basal end of cochlea) are shown as segmented 3D volumes in relation to the round window membrane (RWM), osseous spiral lamina (OSL), limbus (L), and spiral ligament (SL). **c:** A 2D OCT cross-section with labels on the basilar membrane (BM), the inner spiral sulcus (ISS), tunnel of Corti (ToC), space of Nuel (SN), and outer tunnel (OT). **d:** The segmented ISS, ToC, SN, and OT fluid spaces are shown from the basal end of the cochlea with their centerlines visible. The centerlines are used in an ellipse-fitting procedure to estimate the major and minor axes of the cross sections along the length of each volume (e.g. the red arrows shown on the ISS cross section). **e:** Top-down view of the OoC fluid spaces, showcasing the lengths of the segmented volumes. Animal #191.

### 6. Digitizing and 3D-reconstructing histology stacks

The histology slides from the gerbil cochleae were digitized using a Nikon® E800 microscope equipped with a Nikon® DS-Ri2 digital camera. The whole cochlea was imaged with 5× magnification, and with 20× magnification for higher resolution measurements of the OoC hook region. The digitized histology sections were then aligned and segmented in Amira. The whole cochlea was reconstructed in 3D, and the total length of the cochlear spiral was measured by picking out the center point of the OoC region on each image section, and then connecting and fitting the OoC center points with a spline curve along the spiral of the whole cochlea from the basal end to the apex. The OoC centerlines reconstructed from histology and µCT were aligned so that both OoC centerlines started from similar locations near the basal end of the cochlea. Based on this alignment, the distance from the basal end of the cochlea to the location where the OCT imaging was conducted was measured along the OoC centerline. The 3D reconstructions of the OoC fluid spaces from OCT and histology were aligned, analyzed, and compared in Amira. The OCT imaging was with a 10× long working distance lens in the comparable location.

### 7. Table SI-1

The mean (top line in each row) and std (bottom line in each row) of the OoC fluid-space volumes as they varied over time from in vivo to 90-min-PM are listed in Table SI-1.

| Fluid space | Volume normalized by segment length (µL / mm) | | | |
|---|---|---|---|---|
| | OCT (mean±std, N=3) | | | |
| | In vivo | 30-min-PM | 60-min-PM | 90-min-PM |
| ISS | 2.24E-03 | 1.58E-03 | 1.04E-03 | 1.36E-03 |
| | 0.130E-03 | 0.098E-03 | 0.035E-03 | 0.256E-03 |
| ToC | 0.42E-03 | 0.40E-03 | 0.47E-03 | 0.38E-03 |
| | 0.033E-03 | 0.043E-03 | 0.028E-03 | 0.061E-03 |
| SN | 0.31E-03 | 0.50E-03 | 0.46E-03 | 0.35E-03 |
| | 0.048E-03 | 0.009E-03 | 0.090E-03 | 0.148E-03 |
| OT | 0.88E-03 | 0.79E-03 | 0.90E-03 | 1.12E-03 |
| | 0.179E-03 | 0.120E-03 | 0.032E-03 | 0.009E-03 |

**Table SI-1.** Summary of the volumes (per mm) of the OoC fluid spaces for the in vivo and 30-, 60-, and 90-min-PM states, as measured using OCT. All volumes have been normalized by the length of the segmented region, with the values in each row listed in terms of the mean on the first line and std on the line below. This table is related to Fig. 5 in the main manuscript.

### 8. Table SI-2

The mean and std of the ST and OoC-structure volumes as measured using OCT (from in vivo to 90-min-PM) and histology are listed in Table SI-2.

| Volume region | Volume normalized by segment length (µL / mm) | | | | |
|---|---|---|---|---|---|
| | OCT (mean±std, N=3) | | | | Histology (mean±std, N=4) |
| | In vivo | 30-min-PM | 60-min-PM | 90-min-PM | |
| ST | 2.10E-02 | 1.63E-02 | 0.38E-02 | 0.18E-02 | 0.27E-02 |
| | 0.259E-02 | 0.167E-02 | 0.206E-02 | 0.104E-02 | 0.032E-02 |
| OoC structure | 1.17E-02 | 1.41E-02 | 1.45E-02 | 1.54E-02 | 0.42E-02 |
| | 0.199E-02 | 0.160E-02 | 0.117E-02 | 0.297E-02 | 0.16E-02 |

**Table SI-2.** Summary of the ST and OoC-structure volumes (per mm) as measured using OCT (for in vivo through 90-min-PM) and histology. The OCT volumes (N=3) are from single ears of three gerbils, and the histology volumes (N=4) are from both ears of two gerbils (#274 and #338). All volumes are normalized by the length of the segmented region. This table is related to Fig. 7 in the main manuscript.

### 9. Comparing various measures of the OoC cross-sectional area and width

The OoC cross-sectional areas were generated in SimpleWare as part of the centerline analysis. Figure SI-5a compares the mean±se (standard error) of the OoC cross-sectional area as calculated from our in vivo (blue), 90-min-PM (yellow), and histology (green) measurements. The single data point (red triangle) measured by (Richter et al., 2000) is from a hemicochlea preparation on a zero-day postnatal gerbil at a location ~800 µm from the basal end of the cochlea. The in vivo cross-sectional OoC area at this location (18,430 µm$^2$) is less than the 90-min-PM value (21,206 µm$^2$), while the hemicochlea value (2,109 µm$^2$) is much less than either of those values and is about two times smaller than our histology value (5,118 µm$^2$). The hemicochlea data were from a zero-day postnatal gerbil. However, several groups have shown that significant structural and functional changes occur in the gerbil cochlea after birth, and that hearing develops over an approximately three-week postnatal period, during which the dimensions of the OoC structure increase significantly until postnatal day 12, after which the dimensions remain relatively constant (Finck et al., 1972; Müller, 1996; Overstreet and Ruggero, 1998; Souter et al., 1997).

Figure SI-5b compares the mean±se of the OoC_W measurements using the AZ_W+PZ_W method (solid blue and yellow lines) against measurements using the centerline circle-fitting method (dashed blue and yellow lines) for the in vivo (blue; N=5) and 90-min-PM (yellow; N=3) cases. Means are also shown for histology results (red; N=9) from (Plassmann et al., 1987), as well as mean±se results from our own histology (green; N=4). For the AZ_W+PZ_W method, the in vivo (solid blue) and 90-min-PM (solid yellow) cases exhibit overall increases in OoC_W of 112 to 166 µm and 125 to 182 µm, respectively, when moving apically by ~1.1 mm in the hook region. For the circle-fitting method, the in vivo (dashed blue) and 90-min-PM (dashed yellow) cases also show increases in OoC_W, this time of 93 to 190 µm and 141 to 186 µm, respectively, when moving over the same length in the hook region. For the in vivo cases, the two methods yield nearly the same OoC_W values near the 600-µm location, but the circle-fitting method yields a somewhat larger value for OoC_W when moving apically than does the AZ_W+PZ_W method. Because the AZ_W+PZ_W method is based on measuring the widths of cross-sections as defined by the OCT imaging planes, while the circle-fitting method accounts for the 3D curvature of the BM by making the circles orthogonal to the centerline, the AZ_W+PZ_W method likely produces slightly longer widths as the BM curvature increases relative to the plane of the cross-sectional images. However, for the 90-min-PM cases, OoC_W is virtually the same between the two methods. One possible reason for this increase in OoC_W for PM relative to in vivo (except for the in vivo circle-fitting results above 600 µm) could be due to PM cell blebbing and swelling of the OoC structures (Nadol and Burgess, 1985).

The OoC_W measurements from (Plassmann et al., 1987) increase monotonically from 92 µm near the base to 153 µm toward the apex near the 1000-µm location, which generally falls below the in vivo OCT results (blue) and further below the 90-min-PM (yellow) OCT results. Our histological measurements vary from 70 µm near the base to 185 µm near the 1000-µm location. Apical approximately to the 400 um location the OoC_W, from our histology measurements generally has a steeper slope than histology from (Plassmann et al., 1987) and also the OCT measurements. One possibility for this is the change in the cutting angle of the histological sections due to the drastic changes in the curvature that take place in hook region, which needs further investigation. This change in curvature may not be so apparent in the OCT method (circle fitting along the center line vs. AZ_W+PZ_W along imaging planes) because it is likely that the OCT imaging plane is already close to the center line of the BM.

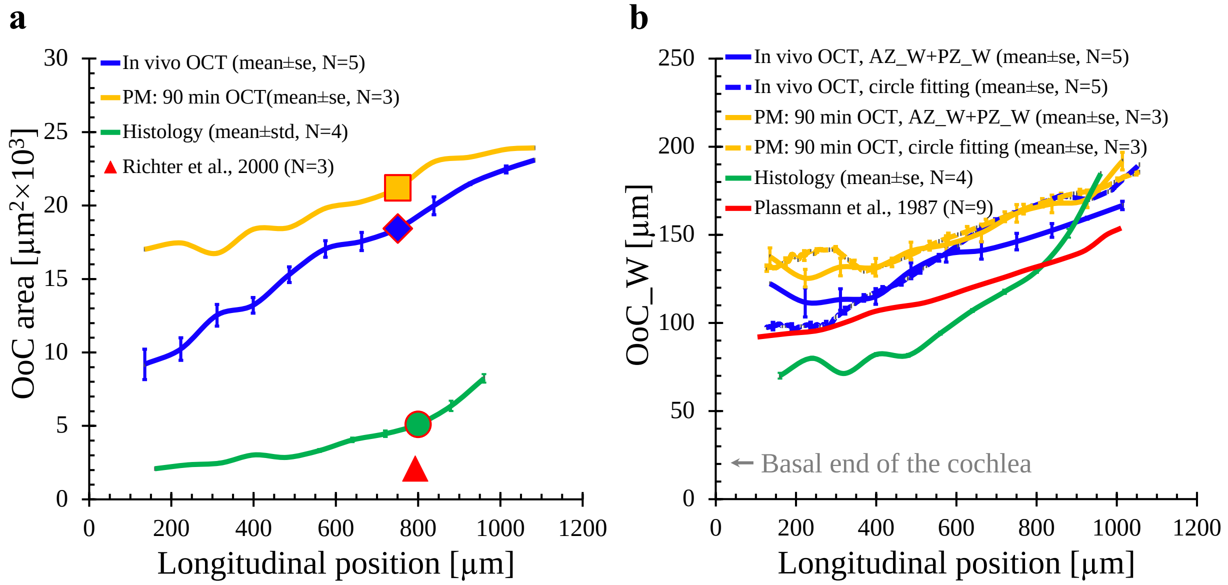


**Fig. SI-5.** Comparisons of OoC cross-sectional areas and OoC_W, as calculated using two methods, from in vivo OCT, 90-min-PM OCT, histology, and measurements found in the literature. **a:** OoC cross-sectional areas are shown for the in vivo (blue; N=5), 90-min-PM (yellow; N=3), and histology (green; N=4) cases, with an added data point at 800 µm from (Richter et al., 2000) (red triangle) based on a hemicochlea preparation from a zero-day postnatal gerbil. To facilitate comparisons, color-matched symbols have been placed at the point closest to 800 µm on the other curves. **b:** OoC_W is shown for the in vivo (blue; N=5), 90-min-PM (yellow; N=3), histology (green; N=4), and histological results (red; mean only; N=9, (Plassmann et al., 1987)). For the in vivo and 90-min-PM cases, OoC_W was calculated using two methods: by adding together the widths of the $BM_{AZ}$ and $BM_{PZ}$ regions based on the OCT-imaging planes (AZ_W+PZ_W; solid lines), and by using the centerline-based circle-fitting method depicted in Figure SI-3 (dashed lines). Aside from the results from the literature, all curves and error bars represent the mean±se (standard error). At some points, the error bars are so short that they are concealed by the line representing the mean.

### References


Finck A, Schneck CD, Hartman AF. 1972. Development of cochlear function in the neonate Mongolian gerbil (Meriones unguiculatus). *Journal of comparative and physiological psychology* **78**:375.

Katsumi S, Sahin MI, Lewis RM, Iyer JS, Landegger LD, Stankovic KM. 2020. Intracochlear Perfusion of Tumor Necrosis Factor-Alpha Induces Sensorineural Hearing Loss and Synaptic Degeneration in Guinea Pigs. *Front Neurol* **10**:1353. doi:10.3389/fneur.2019.01353

Müller M. 1996. The cochlear place-frequency map of the adult and developing Mongolian gerbil. *Hearing research* **94**:148–156.

Nadol JB, Burgess B. 1985. A study of postmortem autolysis in the human organ of corti. *Journal of Comparative Neurology* **237**:333–342. doi:https://doi.org/10.1002/cne.902370305

Overstreet EH, Ruggero MA. 1998. Basilar membrane mechanics at the hook region of the Mongolian gerbil cochlea. *Assoc Res Otolaryngol Mid-Winter Meet Abst* **21**:181.

Plassmann W, Peetz W, Schmidt M. 1987. The Cochlea in Gerbilline Rodents. *BBE* **30**:82–102. doi:10.1159/000118639

Richter C-P, Edge R, He DZ, Dallos P. 2000. Development of the gerbil inner ear observed in the hemicochlea. *Journal of the Association for Research in Otolaryngology* **1**:195–210.

Souter M, Nevill G, Forge A. 1997. Postnatal maturation of the organ of Corti in gerbils: morphology and physiological responses. *Journal of Comparative Neurology* **386**:635–651.
